# Unified neural dynamics of decisions and actions in the cerebral cortex and basal ganglia

**DOI:** 10.1101/2020.10.22.350280

**Authors:** David Thura, Jean-François Cabana, Albert Feghaly, Paul Cisek

**Author notes:** CRNL – ImpAct team, Inserm U1028 – University of Lyon 1, 69675 Bron (France) –. Integrated Regional Center of Cancerology (CRIC), Lévis, QC (Canada). Institute for Research in Immunology and Cancer of the University of Montréal, Montréal, QC (Canada). Corresponding author: Paul Cisek.

## Abstract

Several theoretical models suggest that deciding about actions and executing them are not completely distinct neural mechanisms but instead two modes of an integrated dynamical system. Here, we investigate this proposal by examining how neural activity unfolds during a dynamic decision-making task within the high-dimensional space defined by the activity of cells in monkey dorsal premotor (PMd), primary motor (M1), and dorsolateral prefrontal cortex (dlPFC) as well as the external and internal segments of the globus pallidus (GPe, GPi). Dimensionality reduction shows that the four strongest components of neural activity are functionally interpretable, reflecting a state transition between deliberation and commitment, the transformation of sensory evidence into a choice, and the baseline and slope of the rising urgency to decide. Analysis of the contribution of each population to these components shows differences between regions but no distinct clusters within each region. During deliberation, cortical activity unfolds on a two-dimensional “decision manifold” defined by sensory evidence and urgency, and falls off this manifold at the moment of commitment into a choice-dependent trajectory leading to movement initiation. The structure of the manifold varies between regions: In PMd it is curved, in M1 it is nearly perfectly flat, and in dlPFC it is almost entirely confined to the sensory evidence dimension. In contrast, pallidal activity during deliberation is primarily defined by urgency. We suggest that these findings reveal the distinct dynamics of different regions, supporting a unified recurrent attractor model of action selection and execution.

## Introduction

During natural behavior, we are continuously interacting with a complex and dynamic world^1,2^. That world often does not wait for us to make up our minds about perceptual judgments or optimal choices, and inaction can lead to lost opportunities, or worse. Furthermore, we must often make decisions while we’re already engaged in an action, such as while navigating through our environment or playing a sport^3^. These considerations suggest that the neural mechanisms involved in selecting and executing actions should be closely integrated within a unified sensorimotor control system^4^. Indeed, many neural studies have shown considerable overlap between the brain regions involved in action selection and sensorimotor control^5–10^.

However, while the anatomical overlap between the distributed circuits of decision-making and sensorimotor control is well-established, theoretical models of these processes remain largely separate. Decision-making is often modeled as the accumulation of evidence until a threshold is reached^11–19^, at which time a target is chosen. Models of movement control usually begin with that chosen target, toward which the system is guided through feedback and feedforward mechanisms^20–22^. But if the neural circuits involved in action selection and sensorimotor control are truly as unified as neural data suggests, then theories of these processes should be similarly unified. One promising avenue toward an integrated account of selection and control is to consider both as aspects of a single distributed dynamical system, which transitions from a biased competition between actions^23–28^ into an “attractor” that specifies the initial conditions for implementing the chosen action through feedback control^29–31^.

Here, we test whether neural activity in key cortical and subcortical regions exhibits properties that would be expected from a unified dynamical system for action selection and sensorimotor control. We focus on cells recorded in monkey dorsal premotor (PMd) and primary motor cortex (M1), which are implicated in both selection and control^7,9,32–36^, as well as the dorsolateral prefrontal cortex (dlPFC), which is implicated in representing chosen actions^37^. In addition, we examine activity in the output nuclei of the basal ganglia, the globus pallidus externus (GPe) and internus (GPi), whose role in selection and/or motor control is under vigorous debate^38–41^. Importantly, we examine the activity of all of these regions recorded in the same animals performing the same reach selection task, making it possible to quantitatively compare activities in different brain areas using the same metrics.

To disentangle neural activity related to deliberation, commitment, and movement, we trained monkeys to perform the “tokens task” (Figure 1) (see Methods). In the task, the subject must guess which of two targets will receive the majority of tokens jumping randomly from a central circle every 200ms (Figure 1a). The subject does not have to wait until all tokens have jumped, but can take an early guess, and after a target is reached the remaining tokens jump more quickly (every 150ms or 50ms in separate “Slow” and “Fast” blocks of trials). Thus, subjects are faced with a speed-accuracy trade-off (SAT) – to either wait to be confident about making the correct choice, or to take an early guess and save some time, potentially increasing their overall reward rate. If we assume that commitment occurs shortly before movement onset, then we can delimit within each trial a period of deliberation (Figure 1b) during which neural activity should correlate with the sensory evidence related to token jumps as well as to subjective policies related to the speed-accuracy trade-off. Furthermore, because we can precisely quantify the success probability (SP) associated with each choice after every token jump, we can compute for each trial a temporal profile of the sensory evidence and categorize trials into similarity classes (Figure 1c), including “easy trials”, “ambiguous trials”, and “misleading trials” (see Methods for details).

**Figure 1.**
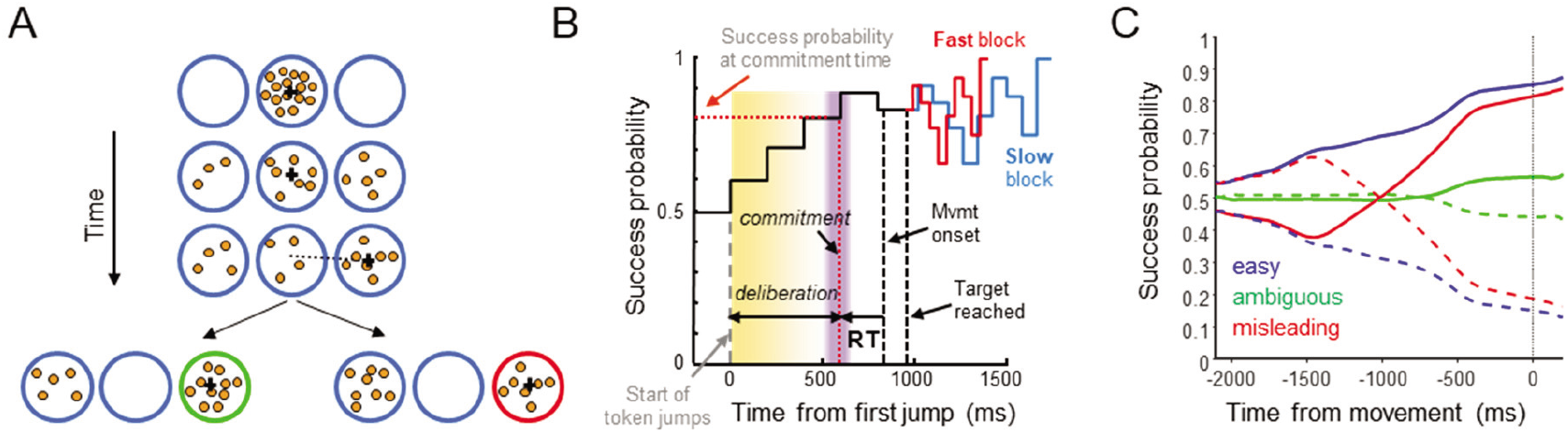
The “tokens task”. **A**. During each trial, 15 tokens jump, one every 200ms, from the central circle to one of two outer target circles. The subject’s task is to move the cursor (black cross) to the target that will ultimately receive the majority of the tokens. **B**. Temporal profile of the “success probability” that a given target is correct. Once a target is reached, the remaining token jumps accelerate to one every 150ms (“Slow” block) or 50ms (“Fast” block). We subtract from movement onset the mean reaction time (RT), measured in a separate delayed-response task, to estimate commitment time (purple bar) and the success probability at commitment time (dotted red horizontal line). **C**. Success probability for choosing the right target in trial types defined on the basis of the success probability profile, here computed after aligning to movement onset (see Methods). Solid: correct target on the right; Dashed: correct target on the left.

Previous studies have shown that both human and monkey behavior in the tokens task is well-explained by the “urgency-gating model” (UGM)^14,42^, which suggests that during deliberation, the sensory “evidence” about each choice (provided by the token distribution) is continuously updated and combined with a non-specific “urgency signal”, which grows over time in a block-dependent manner, and commitment to a given choice is made when the product of these reaches a threshold. Recent neural recordings largely supported these proposals (Extended Data Figure 2). For example, some cells in PMd (29%) and M1 (45%) were significantly tuned during deliberation, reflecting both the evidence favoring their preferred target and the growing urgency signal^36,43^. In contrast, cells in GPe and GPi did not show tuning until after commitment, and about 50% exhibited either increasing or decreasing activity that resembled the context-dependent urgency signal^44^.

While these and other studies reveal important properties of activity in these regions, many questions remain. Are “decision-related” neurons part of a module for choosing a target, which sends its output to a separate module of “movement-related” neurons? Do the basal ganglia contribute to the deliberation process^39,45–49^ or do they simply reflect a choice taken in cortical regions^50,51^ and contribute only to movement execution^52^? Answering these questions is difficult given the heterogeneity of cell properties^7,32,53–56^ and their apparently continuous distribution along rostrocaudal gradients^54,57^ or layers^58^. This leads one to consider whether, instead of serial modules, action selection and execution are two modes of a unified recurrent system distributed across the frontoparietal cerebral cortex and associated basal ganglia/thalamic loops. According to this model^59^, action selection occurs through a competition within the regions of PMd and M1 associated with the relevant effector, biased by signals arriving from dlPFC related to evidence in favor of specific targets. As time passes, that competition is invigorated by an urgency signal coming from the basal ganglia, which gradually amplifies the competitive dynamics in PMd/M1. As the contrast develops between the activity of cortical cell groups associated with different candidate actions, selectivity is gradually induced in the striatum and the pallidum, leading to a positive feedback that further favors the winning cells and suppresses the others, constituting volitional commitment and launching the dynamics of execution^29–31^.

Here, we test this proposal by examining activity across all of the regions we recorded in the tokens task (PMd, M1, GPe, GPi, and dlPFC), but without *a priori* classifying cells into putative functional categories. Instead, we use the neural space approach pioneered in recent years^60–70^, in which the entire system is described as a point in a very high-dimensional space defined by the activity of all recorded cells, and then reduced into a lower-dimensional representation that reveals the main factors governing cell activity across the system. We then perform specific analyses to characterize how activity unfolds during deliberation and commitment, comparing the dynamics of different regions, and quantify to what extent cell properties cluster into distinct functionally interpretable roles. Some of these results have previously appeared in abstract form^71–74^.

## Results

### Neural data

We recorded spiking activity from a total of 736 well-isolated individual neurons in the cerebral cortex and basal ganglia of two monkeys (S and Z), recorded at the locations shown in Extended Data Figure 1. Of these, 356 were recorded in PMd (237 from monkey S), 211 in M1 (79 from monkey S), 62 in dlPFC (60 from monkey S), 51 in GPe (19 from monkey S), and 56 in GPi (22 from monkey S). The properties of some of these neurons have been reported in previous publications, focusing on tuned activity in PMd and M1 and the basal ganglia^36,43,44,75^, as summarized in Extended Data Figure 2. Here, we additionally include neurons recorded in dlPFC, which for technical reasons were almost exclusively obtained in monkey S and to date only described in abstract form^71,72^. While a more complete description of their properties awaits more recordings in additional animals, they are included in the analyses described below, in part to provide a contrast to the other cortical and subcortical areas.

### Dimensionality reduction of neural activity into principal components

To examine how the activity of this entire population evolves over time during the task, we performed Principal Components Analysis (PCA) on all cells recorded in all five brain regions, including any well isolated neuron that was recorded in both Slow and Fast blocks. This included a total of 637 neurons, including 277 in PMd, 191 in M1, 52 in dlPFC, 41 in GPe and 46 in GPi. For the PCA, we used data from only four conditions (right and left choices in the fast and slow blocks) and counted each neuron once. This means that the variance explained was dominated by the PMd and M1 populations, in which we recorded the largest number of neurons. However, reducing the number of cells to 35 in each region did not change the results apart from making them noisier and changing the percentage of variance explained by individual principal components (PCs) (See Extended Data Figure 6a). An alternative approach would be to perform PCA on each population of neurons separately, but this would yield region-specific PCs that make quantitative comparisons between regions impossible. Thus, we elected to perform PCA on all cells together, counting each neuron once, producing a “loading matrix” of coefficients (from each neuron to each PC) that then allows us to “project” each population into the space of the same PCs. As shown below, this allows us to directly compare the components of different regions and to infer how each region contributes to the same distributed dynamical system. We imposed symmetry on our population by following the “anti-neuron” assumption, which assumes that for every neuron we recorded there exists a similar neuron with the opposite relationship to target direction (even for cells that are not tuned), effectively doubling the number of neurons. See the Methods section for the justification and motivation for this approach.

The first 20 PCs together explained 97.9% of the variance in activity over time across the four groups of trials (slow/fast blocks x left/right choices) used for the Principal Components Analysis. Figure 2 shows the temporal profile for the first 7 PCs constructed as a weighted average of all cells, separately for easy, ambiguous, and misleading trials in both slow and fast blocks, for both left and right choices. Here, the PCs are calculated after aligning the data to movement onset. See Extended Data Figure 3 for the same PCs calculated after aligning to the beginning of the trial.

**Figure 2.**
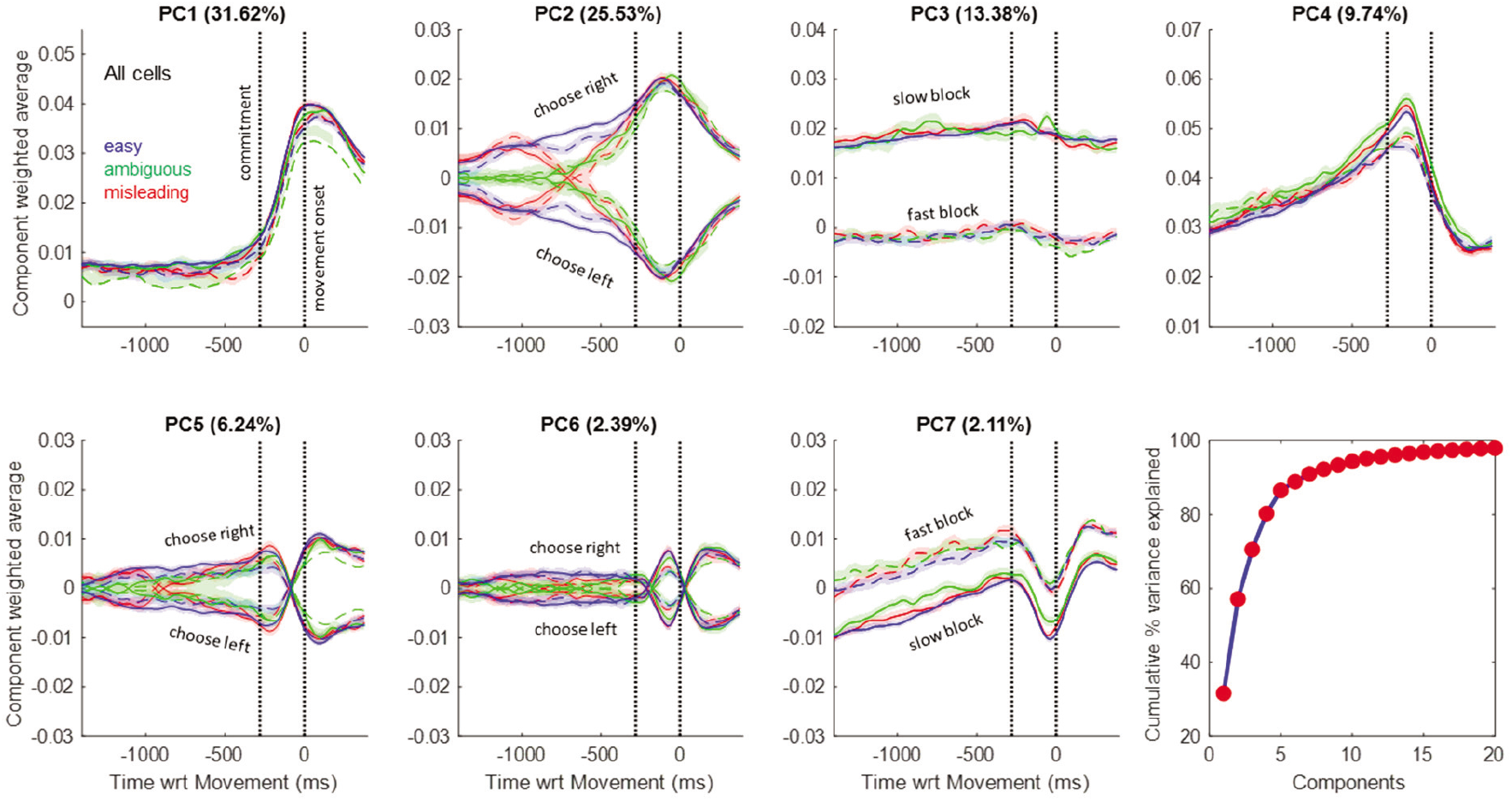
Components produced by PCA. The cumulative variance explained by the first 20 components is shown at the bottom right, and the temporal profiles of the top 7 PCs are shown in the rest of the figure. Each of those 7 panels shows the average activity of all cells, weighted by their loading coefficient onto the given PC, for 12 trial groups, including combinations of 3 trial types: easy (blue), ambiguous (green), and misleading (red); two blocks: Slow (solid) and Fast (dashed); for choices made to the left or right (indicated for the components where they differ). Note that the sign is arbitrary because the loading matrix can have positive or negative values. Shaded regions indicate 95% confidence intervals. In each panel, the second vertical dotted line indicates movement onset and the first indicates the estimated moment of commitment, 280ms earlier, based on our prior studies. Each panel is scaled to have the same range in the y-axis. PCs for the Slow trials are built from 557 cells that have all trial types in Slow blocks, while PCs for Fast trials are built from 452 cells that have all trial types in Fast blocks.

The first 4 components, which together explain 80.3% of total variance, are clearly interpretable in terms of the key elements of the urgency gating model. The first PC (31.62% of variance) is nearly identical across all conditions and reflects the transition between deliberation (prior to commitment) and action (after movement onset). It is similar to the main condition-independent component reported in other neural space studies in both primates^76–80^ and rodents^81–83^. The second PC (25.53%) exhibits two phases. Prior to commitment, it reflects the time course of the sensory evidence on which the monkey made his choice, distinguishing easy, ambiguous, and misleading trials. After commitment, it simply reflects the choice made without distinguishing trial types. The third PC (13.38%) reflects the block-dependent aspect of urgency, distinguishing between the slow and fast blocks (even before the start of the trial, as shown in Extended Data Figure 3). The fourth PC (9.74%) reflects the time-dependent aspect of urgency until just before movement onset. The remaining components are similar to PCs 2 and 4 during deliberation, but capture some of the heterogeneity across cell activity patterns after commitment and movement onset. We discuss these higher PCs in the Supplemental Materials.

PC2 warrants special attention. Note that until the moment of commitment, it correlates very well with the evidence provided by the token movements (compare to Figure 1c). Indeed, as shown in Extended Data Figure 4, the correlation between the time-delayed evidence and the value of PC2 is highly significant (p<10^-100^) with a correlation coefficient of R=0.92. This is notable because the PCA algorithm was not given any information about these different trial types (easy, ambiguous, misleading, etc.) but was simply given data averaged across four large trial groups that only distinguished left versus right choices and slow versus fast blocks. Nevertheless, when the resulting temporal profiles of PC2 are calculated for specific trial types they clearly reflect how the evidence dynamically changes over the course of deliberation in those trials.

Could this finding be a trivial consequence of overall cell tuning? To test this possibility, we used the Tensor Maximum Entropy method of Elsayed & Cunningham^84^ to generate synthetic data sets that retain primary features such as tuning, but are otherwise random (see Methods). As shown in Extended Data Figure 4c, when PCA is applied to such synthetic data sets it does not produce components that are as well correlated with evidence as PC2 from our true data (p<0.01). The implication is that the emergence of PC2 in the real neural data requires a consistent relationship between how cells reflect the final choice (left vs. right) and how they reflect the evidence that leads to that choice during deliberation (easy vs. ambiguous vs. misleading, etc.).

Figure 3 shows the trajectories of the different trial types, separately for the Slow and Fast blocks, plotted in the space of PCs 1, 2, and 4. While a quantitative comparison between the Slow and Fast blocks is made difficult because a slightly different subset of cells is included in each (see Methods), the qualitative shape of the trajectories is very similar. As indicated in Figure 3a (dotted black arrows), in both block types the neural activity evolves in a clockwise manner in the space of PC1 and PC4, passing over a region of deliberation until reaching a commitment state (purple ellipses), whereupon it rapidly moves to a movement-specific initiation subspace (green ellipses), and then turns back toward the starting point during movement execution. Some of these phenomena have previously been reported using neural space analyses of preparatory and movement-related activity in cortical regions during instructed reaching tasks with a single target^76–79,85–87^. Here, our task allows us to examine in more detail what happens during the process of prolonged deliberation when subjects are selecting among multiple targets.

**Figure 3.**
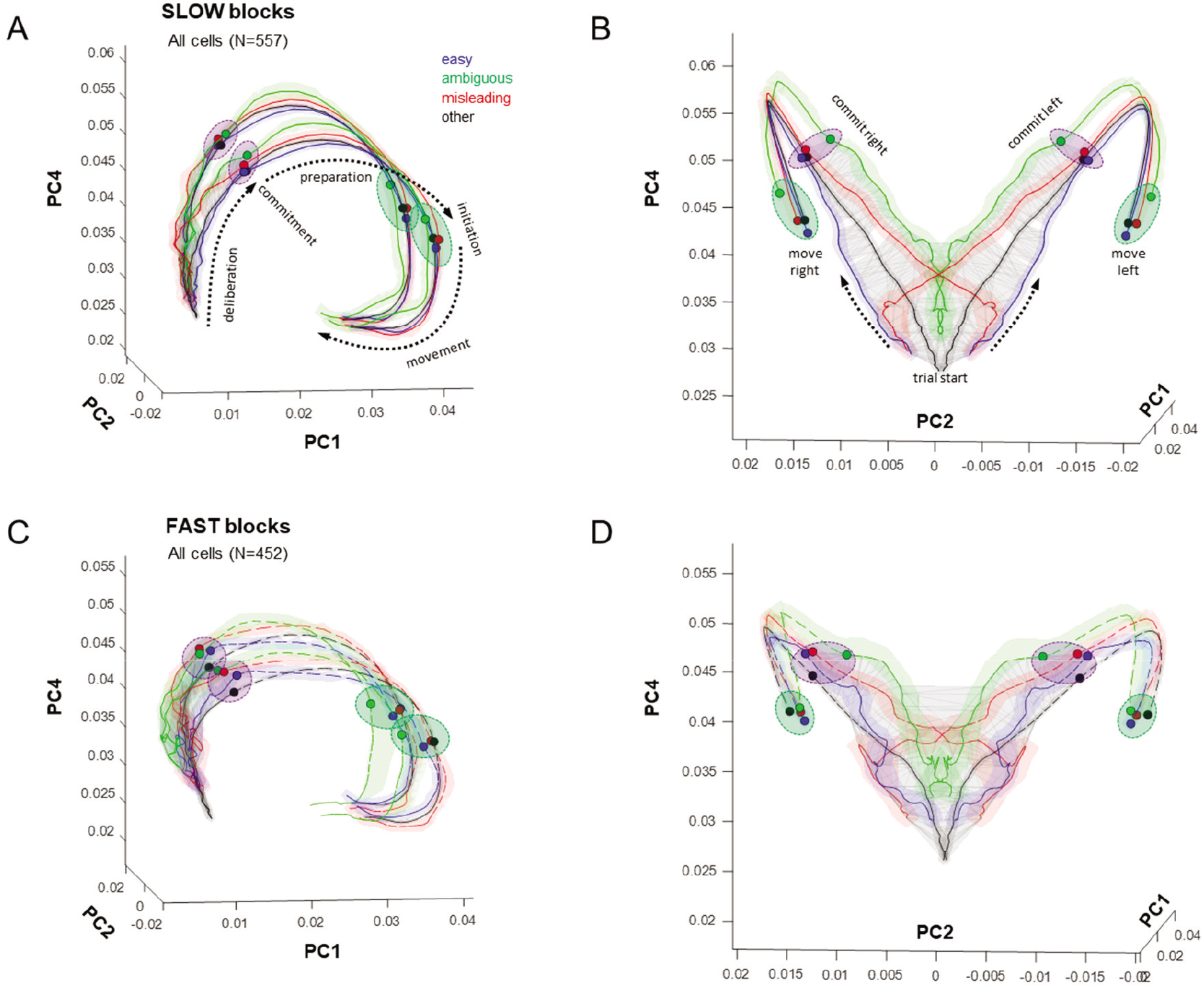
Neural trajectories in the space of PCs 1, 2 and 4, averaged across easy, ambiguous, and misleading trials as well as the other (unclassified) trials. Panels A and B show two views of data from Slow blocks, and panels C and D show the same views of data from the Fast blocks. Shaded colored regions around each trajectory indicate the 95% confidence interval. Dotted arrows in A indicate how the state of activity evolves over time. The gray wireframe encloses all states visited during the deliberation epoch, across all trial types. Purple ellipses indicate the region in which commitment occurs (indicated for individual trial types by small colored circles) and green ellipses indicate the point at which movement is initiated. For clarity, the neural state after movement initiation is only shown in panels A and C.

In each panel of Figure 3, we’ve drawn a gray wireframe around all of the points from the beginning of the trial until commitment time (280ms before movement onset), across all trial types, thus defining the subspace within which deliberation occurs. This subspace resembles a triangular surface that is extended mostly in PC2 and PC4 and curved slightly into PC1. It is quite thin – for example, in the slow block the value of Ψ (see Methods) is 0.204, which is roughly equivalent to a triangular sheet whose thickness is 1/57^th^ of the length of each side. We call this the “decision manifold”.

The flow of neural states upon the decision manifold is quite orderly, proceeding from bottom to top as time elapses and shifting left and right with the sensory evidence. For example, consider the misleading trials, in red, which clearly reveal the switch in sensory evidence. In effect, the flow of the neural state during deliberation resembles the temporal profile of evidence (Figure 1c) mapped onto that curved wireframe surface. The neural state continues to flow along the decision manifold until it reaches one of two edges (purple ellipses) at the time of commitment, and then turns into PC1 and accelerates to rapidly flow along one of two paths, each corresponding to the choice taken, until movement initiation (green ellipses).

### Region-specific dynamics

While the structure of the neural space computed across all neurons is interesting, it is still more informative to compare that structure across the different brain regions in which we recorded. Because the loading matrix produced by PCA provides coefficients that map each individual neuron’s contribution to each PC, we can “project” the activity of any subset of neurons into the space of these same PCs (see Extended Data Figure 5), and then plot region-specific neural space trajectories. Figure 4a shows this for the dorsal premotor cortex, where we see a structure that is quite similar to what was shown for all neurons (not surprisingly since the PMd population is the largest). As before, we see a triangular decision manifold that is relatively thin (Ψ=0.343) and extends along PC2 and PC4. However, note that it initially strongly leans in the negative PC1 direction and then curves around just before commitment (see the side view shown in Figure 4a, right). Interestingly, the PMd decision manifold is curved as if it lies on the surface of a sphere (see inset, spherical fit R^2^ = 0.65), a point to which we will return below.

**Figure 4.**
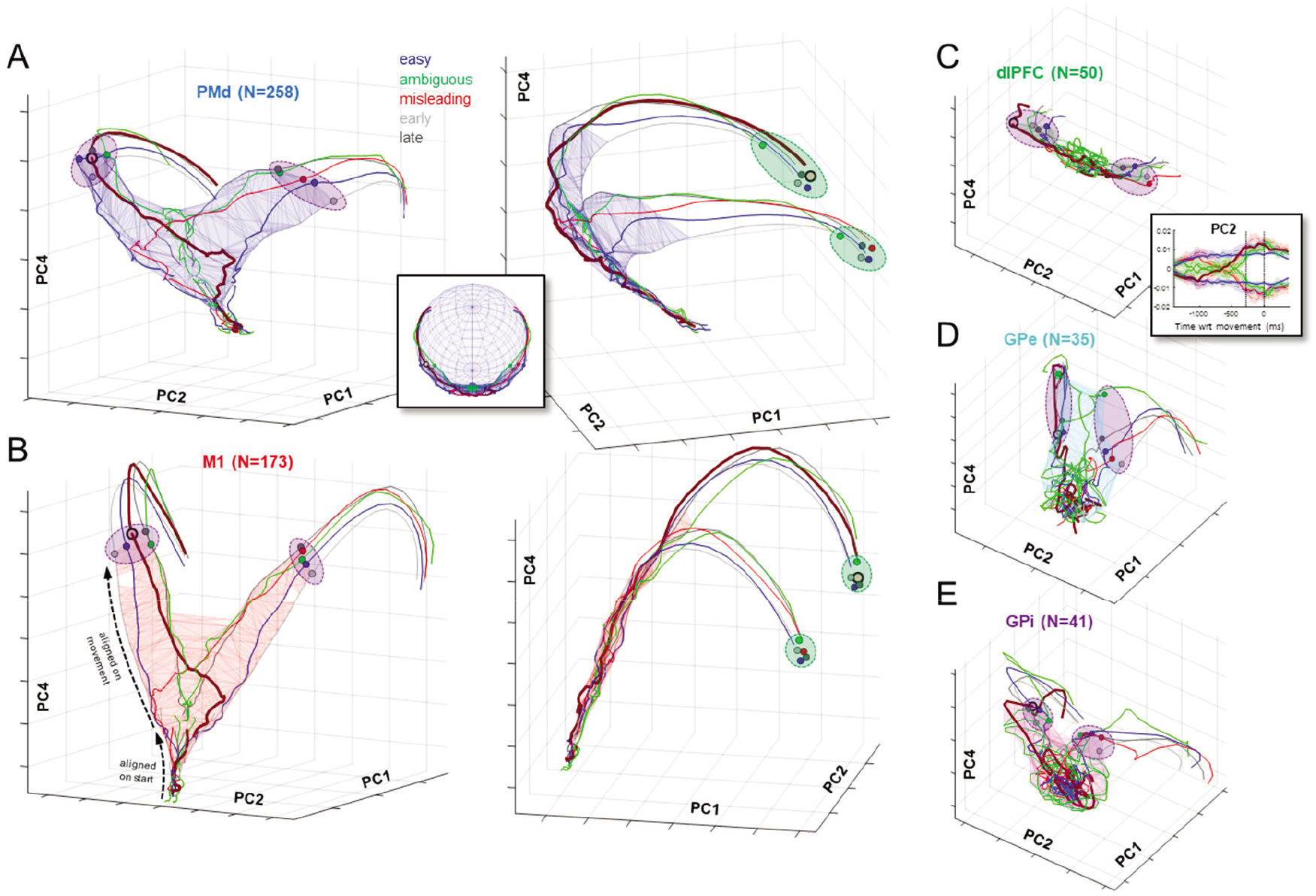
Neural trajectories in the space of PCs 1, 2 and 4, plotted using activity during Slow blocks, separately for PMd (A), M1 (B), dlPFC (C), GPe (D) and GPi (E). Separate trajectories are plotted for easy, ambiguous, and misleading trials, as well as all trials in which decisions were shorter than 1400ms (early) and in which decisions were longer than 1400ms (late). For clarity, confidence intervals have been omitted. As in Figure 3, trajectories are computed from data aligned on movement onset, and extend from 1400ms prior. Blue (PMd) and red (M1) wireframes enclose all states before commitment (280ms before movement). Here, we also superimpose trajectories computed from data aligned on the start of the trial and until 500ms later (projected through the same loading matrix). Dashed black arrows in panel B, left, indicate where these separate trajectories can be seen in the M1 space. In all panels, the trajectory of misleading trials in which the monkey correctly chose the right target is highlighted with a thicker line. Purple ellipses emphasize the time of commitment and green ellipses emphasize movement onset. The inset in A shows how PMd activity tends to remain on the surface of a sphere, particularly during deliberation. The inset in C shows PC2 computed from the dlPFC population for easy (blue), ambiguous (green), and misleading trials (red).

In contrast, the decision manifold of primary motor cortex (M1, Figure 4b) is remarkably flat and thin (Ψ=0.252) and leans into the positive PC1 direction. Nevertheless, the evolution of the neural state along the surface of the decision manifold in both regions obeys the same pattern seen in Figure 3, proceeding from bottom to top as time elapses and shifting left and right with the sensory evidence, always lying within the same subspace (curved for PMd, planar for M1). Note that Figure 4 shows data aligned on movement onset, as in previous figures, but also superimposes neural space trajectories computed on the basis of data aligned on the start of the trial (see Extended Data Figure 3) and then projected into the same PC space using the same loading matrix. As is clear from the figures, regardless of how the trajectories are computed, they always fall within the same decision manifold in PC space, for both PMd and M1.

It is noteworthy that in both PMd and M1, the state reached at the moment of commitment (purple ellipses in Figure 4a,b) shows an orderly relationships with the reaction time in each trial type (shortest in “early” trials, and longest in “late” trials). This is in agreement with the observation that even at a single trial level, a consistent relationship between neural state and reaction time can be observed during a simple instructed reach task^76^. Furthermore, in M1 the trajectories after that point converge to arrive in a relatively compact subspace (green ellipses) at movement onset - what Churchland et al.^88^ called an “optimal subspace”.

In contrast to PMd and M1, the deliberation manifold in dorsolateral prefrontal cortex (dlPFC, Figure 4c) is almost exclusively extended along PC2 (Ψ=0.564, like a cylinder whose length is 22 times its radius). Like PMd and M1, the neural state along PC2 shifts left and right with sensory evidence (see inset), but after commitment it exhibits only a small excursion into PC1. This suggests that neural activity in dlPFC primarily reflects the sensory evidence used to make decisions in the task, consistent with many previous studies^89–96^.

A strong contrast to the cortical data is seen when we examine the neural state of cells recorded in the globus pallidus (GPe and GPi). In both regions, the neural state during deliberation is confined within a subspace that is not a thin manifold but instead resembles a ball compressed along PC2 and extended along PC4 (Figure 4d,e). During deliberation, activity in these regions does not evolve in an orderly fashion as seen in cortex. This could be partly due to the lower number of cells recorded in GPe and GPi as compared to PMd and M1, although similar results hold when we restrict *all* regions to 35 cells (Extended Data Figure 6a). Nevertheless, by the time commitment occurs, the state of both GPe and GPi lies in a choice-specific subspace (purple ellipses) and then evolves quickly to a corresponding initiation subspace. These findings are consistent with our previous report of GPe/GPi activity in the tokens task, in which we suggested that these regions do not determine the choice but rather contribute to the process of commitment^44^.

### Interpreting the shape of decision manifolds

As shown in Figure 4, the decision manifolds computed from PMd and M1 possess distinct shapes. While M1 is almost completely flat, the PMd manifold is curved, as if lying on a surface of a sphere. Do these shapes reveal differences in the neural dynamics in these regions? To address this question, we consider the shape of the PMd decision manifold in terms of two separate phenomena: its curvature in the PC1-PC4 plane (Figure 4a, right), and its curvature in the PC1-PC2 plane (inset in Figure 4a).

First, we consider why the PMd manifold initially leans in the negative PC1 direction and then bends toward the positive PC1 direction prior to commitment (its curvature in the PC1-PC4 plane). One potential explanation suggests that an inhibitory influence prevents premature movement before selection is complete^97^ by keeping PMd away from commitment, but it is gradually overcome by positive feedback in the recurrent circuit between PMd and GPi^44,59,98^. According to this hypothesis, as the cortical activity becomes increasingly biased in favor of one target over another, it gradually begins to produce the emergence of choice selectivity in the GPe, about 200ms before commitment (see Extended Data Figure 2b, row 3). When that becomes strong enough to engage selectivity in the GPi, it in turn strengthens emerging selectivity in the thalamus, which then further strengthens selectivity in the cortex. Thus, a positive feedback is established leading to a winner-take-all process that overcomes inhibition and constitutes volitional commitment. This hypothesis predicts a relationship between how tuning emerges in GPi and how and when the PMd state begins to flow toward commitment.

To test this prediction, we examined the correlation between the flow toward commitment in PMd (as reflected in the rate of change of PC1 during deliberation) with the depth of directional selectivity in GPi (as reflected in the absolute value of PC2). Figure 5a shows this comparison for 6 large trial groups: all rightward choices in Slow and Fast blocks, early rightward choices in Slow and Fast blocks, and late rightward choices in Slow and Fast blocks (see Methods for trial definitions). Leftward decision trials were not considered because they are symmetric with rightward decision trials and thus redundant. As can be seen for the different trial types, except for a constant scaling factor the match between these two very different variables is strong, particularly in the epoch between commitment and movement onset. Even if the data is restricted to the period *before* commitment, the correlation for each trial group shown in the individual panels of Figure 5a is significant at p<10^-10^ with Pearson’s R above 0.86, and for all trials together with R=0.84 (Figure 5b). Furthermore, across these six conditions there was a significant correlation (p=0.0029, R=0.9556) between the time when the PMd manifold began to tilt toward commitment (derivative of PC1 became consistently positive) and the time when tuning became significant in GPi (95% CI of PC2 no longer included 0). Similar results were not obtained when comparing GPi PC2 against the derivative of PC1 computed from other regions (although there was a weaker but significant correlation with the derivative of PC1 in M1). It is important to note that this particular prediction – a relationship between the derivative of one component in PMd and the absolute value of another in GPi – is not arbitrary. It is motivated by the specific proposal that the timing of the bend in the PMd decision manifold in the PC1-PC4 plane (Figure 4a, right) is related to the timing of how tuning emerges in the basal ganglia, which is itself motivated by the hypothesis that both of these variables reflect positive feedback in a recurrent attractor circuit. Of course, like any correlation analysis, it cannot conclusively prove a specific causal relationship. An alternative, though not mutually exclusive hypothesis is that both of these regions are influenced by a common source of inhibition that is gradually released as commitment approaches.

**Figure 5.**
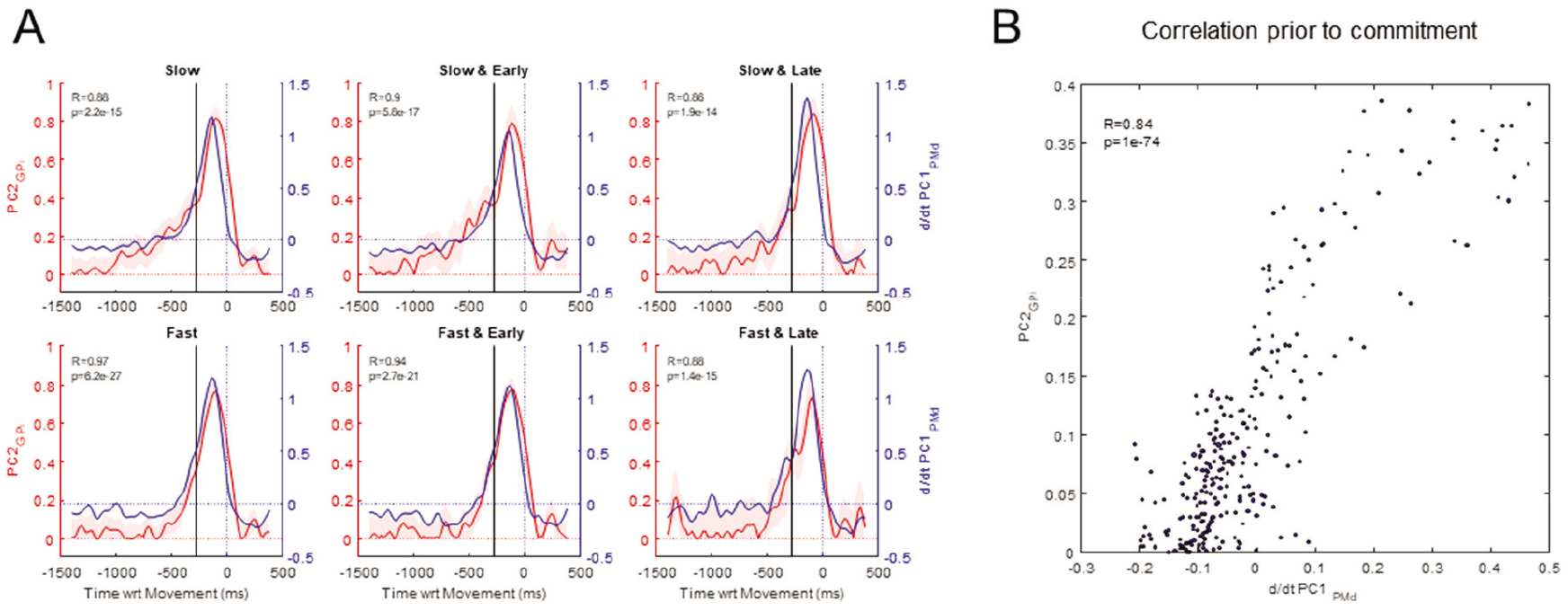
Comparison of selectivity in GPi with the slope of the PMd decision manifold. **A**. Comparison within six conditions: all Slow block trials, Slow block trials where DT<1400, Slow block trials where DT>1400, all Fast block trials, Fast block trials where DT<950, Fast block trials where DT>950. Each panel plots the absolute value of PC2 computed from GPi (red, left y-axis, shaded region indicates 95% CI) along with the derivative of PC1 computed from PMd (blue, right y-axis). The black vertical line indicates the estimated time of commitment. At the top left corner are Pearson’s R and the p-value of the correlation between these two signals from 1000ms to 280s before movement. **B**. The correlation of these signals, from 1000 to 280ms before movement onset, across all 6 conditions.

In addition to its curvature in the PC1-PC4 plane, the PMd manifold is also curved in the PC1-PC2 plane, as if it lies upon a surface of a sphere (see inset in Figure 4a). This is strikingly different than the M1 manifold, which is nearly perfectly flat (compare Figure 4a with b). What could explain this difference in shapes?

Here, we consider one possible explanation related to the dynamics of recurrent attractor networks. We illustrate this using a very simple system consisting of two neurons that compete against each other through recurrent inhibition. Note that this minimal model is not intended to simulate our data, but simply to demonstrate the possibility that some features of our data (e.g. shape of the decision manifold) could be the result of very general properties of recurrent non-linear dynamical systems.

Let’s consider two neurons whose activity is denoted as *x*_1_ and *x*_2_ (Figure 6a) and governed by the following differential equation^23^

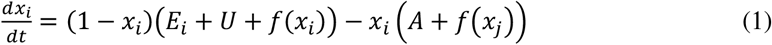

On the right-hand side of equation (1), the first term is excitation and the second is inhibition. *E_i_* is the input evidence for choice *i, U* is the urgency signal, *A* is a passive decay rate, *f*(*x_i_*) is the recurrent excitation of each cell to itself, and *f*(*x_j_*) is the recurrent inhibition from the other cell, *j*. Note that equation (1) ensures that cell activities are always in the interval from 0 to 1. The function *f*(*x*) is sigmoidal of the form

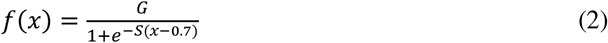

where *G* is the gain and *S* is the steepness of the sigmoid. Figure 6 schematizes this simple model and illustrates some of its dynamics.

**Figure 6.**
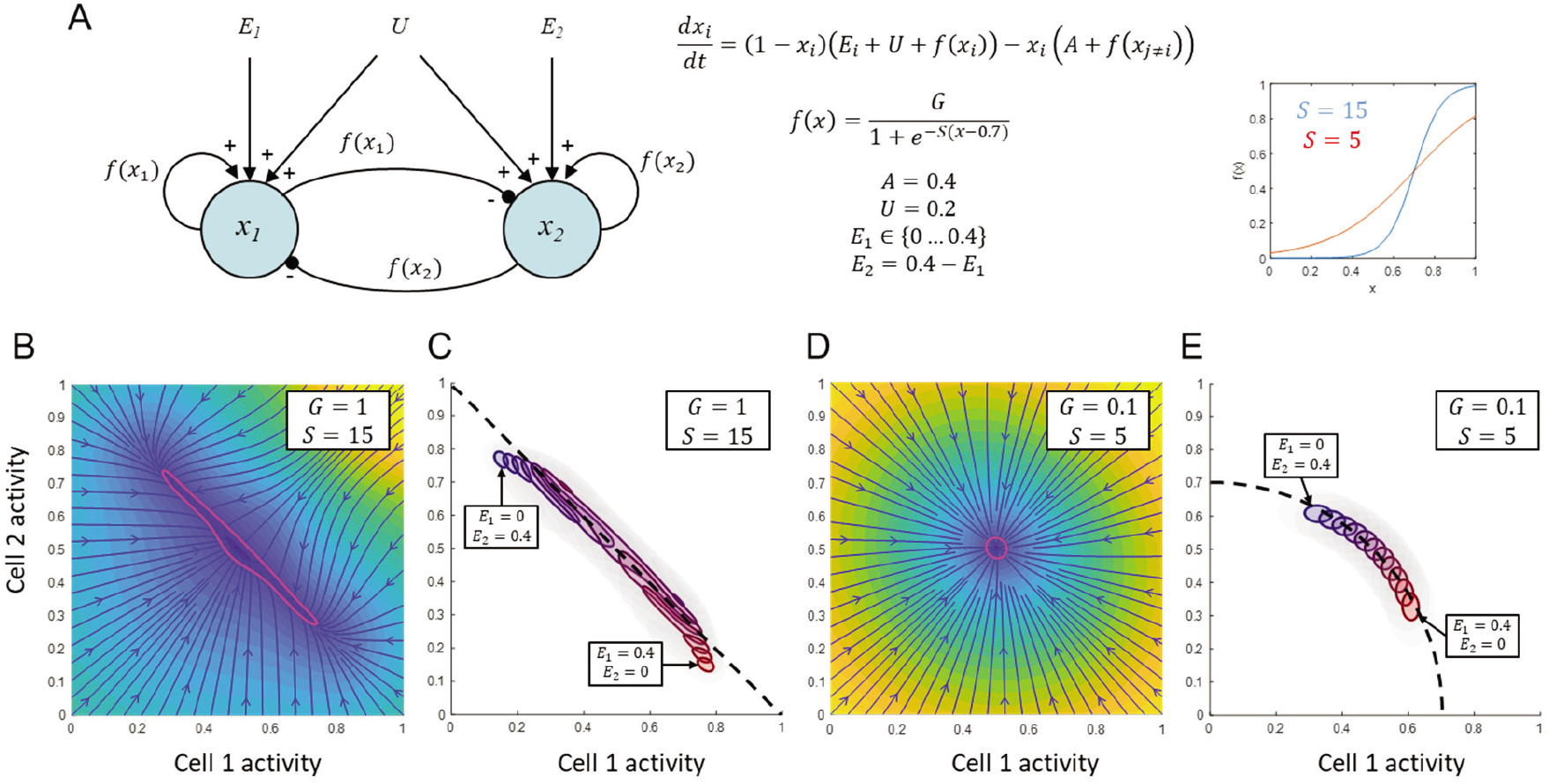
Simulation of a simple 2-neuron recurrent competitive attractor. **A**. The structure of the model is shown at left, next to the governing equations and parameter settings. Two forms of the interaction function *f*(*x*) are shown in the inset at right. **B**. The flow field for a system with high gain (G=1) and steep slope (S=15) when the input evidence is balanced such that E_1_=E_2_=0.2. Blue arrows depict the flow and shading indicates the speed (yellow=fastest, dark blue=slowest). The purple outline indicates a region around the stable equilibrium, in which the state will tend to remain even with substantial noise. Note that the dynamics flow quickly toward that region and then move more slowly within it. **C**. The stable regions of the same system for several input patterns, ranging from strong evidence in favor of choice 2 (top left, blue) to strong evidence in favor of choice 1 (bottom right, red). Note that these align in what is approximately a straight line across the state space (compare to dashed black line). **D**. Same as B but for a system in which the interaction function has low gain (G=0.1) and a shallow slope (S=5). **E**. Stable regions of the system in D. Note that now, the stable regions for the same range of input patterns do not form a straight line but rather fall on a curve roughly equivalent to a circle (dashed black curve).

Figure 6b shows the flow field of the system in the (*x*_1_, *x*_2_)-plane assuming a function *f*(*x*) with high gain and a steep slope, when the evidence is balanced. Note how the flow field quickly pushes the neural state to a central stable region, outlined in purple, within which the flow is slower. In the presence of noise, the system will be strongly constrained to remain inside this region, but can shift within it relatively easily. As the balance of evidence changes, the stable region shifts in the plane (Figure 6c) but always lies oriented along that straight line. As a result, activity is normalized such that *x*_1_+*x*_2_ is approximately constant (L1-normalization). In contrast, Figure 6d,e shows a system where the interaction function has low gain and a shallow slope. Note that the stable region for different input patterns still shifts in the (*x*_1_, *x*_2_)-plane, but not along a straight line (panel e). Instead, the different stable regions now lie along a circular path, such that 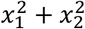 is approximately constant (L2-normalization). In both systems, the state is strongly confined into a narrow subregion of the full space (a “decision manifold”), but can shift within it due to changes of evidence as well as noise. If the value of *U* is increased, this subspace will shift toward the upper right (not shown) until the system bifurcates into two stable attractors, each corresponding to choosing either *x*_1_ or *x*_2_.

This simple model proposes a straightforward candidate explanation for the difference in shapes of the decision manifolds in PMd versus M1 (Figure 4a,b). To summarize, relatively gentle competitive dynamics between candidate options can produce a curved quasi-spherical manifold, as seen in PMd, while strong and steep winner-take-all dynamics can produce a flat one, as seen in M1. This may imply that PMd tolerates multiple competing potential actions more readily than does M1, a conjecture made in previous modeling work^25^. However, at present these are just conjectures. A more complete analysis would require a more sophisticated model, in which different neural populations are dynamically coupled and interact in more complex ways. It is possible that in such a model, other parameter settings and other features of dynamics may better explain the shapes of decision manifolds, but exploring those possibilities is beyond the scope of the present paper.

### Analyses of the loading matrix

Another approach for inferring the putative functional contributions of different brain regions is to examine the distribution of the loading coefficients for cells in each region. For example, if a given population of cells is strongly related to the sensory evidence provided by token jumps, then cells in that population should tend to have higher loading onto PC2 than cells from another population that is less sensitive to evidence. As described in Methods, we characterized the distribution of loading coefficients for the first 11 PCs (which capture 95% of variance) by fitting the 11-dimensional space of points for each brain region with gaussian mixture models (GMMs). The results are shown in Figure 7.

**Figure 7.**
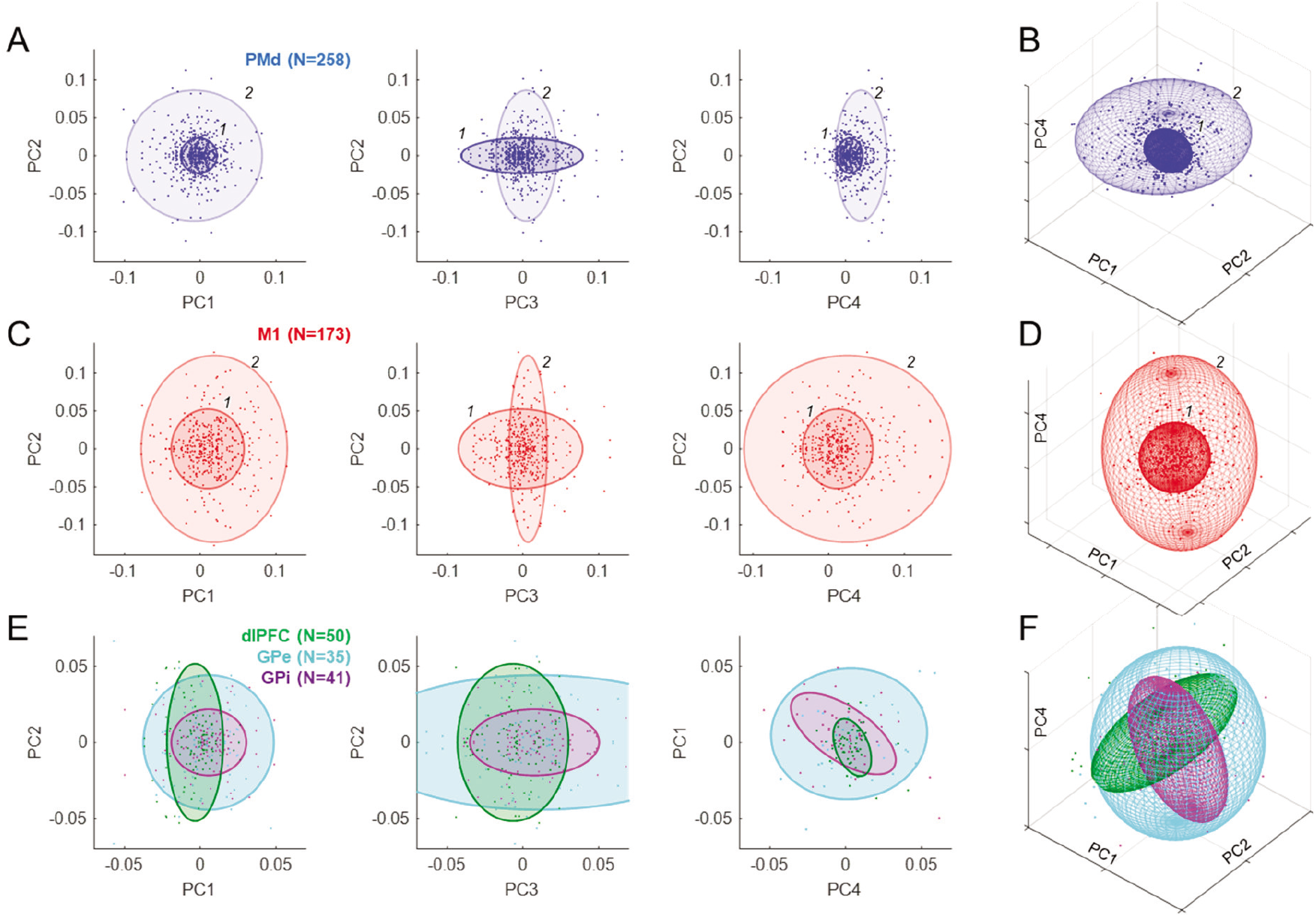
Analysis of loading coefficients. **A**. Each point indicates the weight of the contribution of a given PMd cell to the principal component indicated on the axes. Colored ellipses indicate the centroid and 3 times standard deviation of each of the two Gaussians (labeled *1* and *2*) that provided the best fit to the distribution of these populations. **B**. The same PMd Gaussians in the 3-d space of PC1, PC2, and PC4. **C**. Same as A for M1. **D**. Same as B for M1. **E**. Same as A for cell populations in dlPFC (green), GPe (cyan), and GPi (purple), each of which was best fit with a single Gaussian. Note that unlike A and C, here the right panel shows PC4 vs PC1. **F**. Same as B for dlPFC, GPe, and GPi.

The distribution of loading coefficients for cells in PMd (Figure 7a) was highly distributed but not without structure, and was best fit with two gaussians. The first (*1*: 68% contribution to the fit) was dominated by cells only weakly contributing to PCs 1, 2, and 4, but strongly to PC3. The second (*2*:32%) included cells that strongly contributed to PCs 1 and 2 but more weakly to PCs 3 and 4. The distribution for M1 was also best fit with two gaussians, one (*1*: 56%) contributing mostly to PC3 and another (*2*: 44%) contributing to 1, 2, and 4 but not 3. Thus, in both PMd and M1, there was a trend for cells that most strongly reflect the animal’s speed-accuracy trade-off (PC3) to be less strongly tuned to direction (PC2), and vice-versa (Figure 7a,c, middle panels).

In contrast, populations in dlPFC, GPe, and GPi were well fit with a single gaussian (Figure 7e,f), though perhaps more structure would have been seen with larger populations. Importantly, there were significant and potentially functionally relevant differences between these regions. In particular, the distribution of loadings from dlPFC and GPi were nearly orthogonal in the space of PCs 1, 2, and 4 (Figure 7f). The dlPFC was extended along PC2 and not along the other components, consistent with the proposal that it primarily carries information on the sensory evidence provided by token movements. In contrast, GPi was relatively narrow in PC2 and instead extended along PCs 3 and 4, related to the block and time-dependent aspects of urgency. Furthermore, there was a significant negative correlation (R=-0.32, p<0.001) between GPi loadings on PC1 versus PC4 (Figure 7e, right panel, purple). In other words, cells that build-up over time (positive on PC4) tend to reduce their activity after commitment (negative on PC1), while cells that decrease over time (negative on PC4) tend to increase after commitment (positive on PC1).

In summary, our analyses of loading matrices suggest that while different regions do appear to contribute to different subsets of PCs (e.g. the orthogonal relationship of dlPFC and GPi in Figure 7f), the population within each region does not contain distinct clusters. In other words, while there is some structure in the loading matrix (e.g. Figure 7a,b, middle panel), the distributions of properties within each region are continuous.

Could that continuity of properties be an artifact resulting from dimensionality reduction? That is, if distinct categories of cells with different functional roles really did exist, would our analyses be able to identify them? To address this question, we created a variety of synthetic populations of neuron-like units and applied to them the same analyses we used to examine real data, including PCA and GMM analyses of the resulting loading matrix. As described in the Supplemental Materials (see Extended Data Figure 8), this yielded three conclusions: It confirmed that PCA does correctly identify all of the components from which our synthetic neural populations were constructed. However, it also showed that some of the higher-order components (like PC5 in Figure 2) can result from PCA “cancelling out” some of the firing patterns already captured by lower-order components (e.g. PC2), in order to explain things like neurons tuned only during movement. Nevertheless, in all cases the GMM analysis of the loading matrix correctly identified any real categories of neurons in the population. This suggests that the lack of distinct clusters we found in our data indeed reflects the absence of separate categories in the real neural populations.

## Discussion

Many neurophysiological studies have suggested that decisions between actions unfold within the same sensorimotor regions responsible for the control of those actions^1,99–101^. This includes FEF and LIP for gaze choices^6,91,102–105^ and PMd and MIP for reaching choices^7,36,105–107^. Several computational models of the decision mechanism suggest that it behaves like a “recurrent attractor system”^23–28^, where reciprocally competing groups of neurons tuned for the available choices compete against each other until one group wins and the system falls into an attractor corresponding to a specific choice.

The results reported here provide strong support for this class of models. In particular, the cells we recorded in PMd and M1 do not appear to belong to separate categories related to decision-making versus movement preparation or execution, but instead behave like part of a unified dynamical system that implements a biased competition and transitions to commit to a choice through a winner-take-all process. During deliberation, the pattern of cell activity in these regions is confined to a highly constrained subspace in the shape of a thin manifold, and is shifted around within that manifold by the decision variables pertinent to the task (here, the sensory evidence and the rising urgency). When commitment occurs, the same group of cells now transitions from the decision manifold to a roughly orthogonal tube-shaped subspace corresponding to a specific choice^81^ and quickly flows to a subspace related to movement initiation^77,88^. This is precisely the kind of “winner-take-all” phase transition that occurs in recurrent attractor models.

Additional insights can be obtained by examining the low-dimensional components produced by dimensionality reduction. In particular, it is noteworthy that during deliberation, the four strongest components of neural activity (which together account for just over 80% of the variance) capture the key elements of the urgency-gating model^14^: The momentary evidence (in PC2), the urgency signal (a context-dependent baseline in PC3 and time-dependent ramping in PC4), and the transition to commitment (PC1). Perhaps even more important is how these components are differently expressed in the different cell populations. In particular, the neural state in dlPFC varies almost entirely along PC2 while, conversely, the state in GPe/GPi during deliberation is primarily determined by PC3 and PC4, and not at all by PC2. This suggests that information about sensory evidence is provided by prefrontal cortex^89–96^ while the urgency signal is coming from the basal ganglia^44,108,109^.

The presence of the evidence-related component PC2 in the cortical data is particularly remarkable because the dimensionality reduction algorithm was not provided with any information about the variety of trial types (easy, ambiguous, etc.) but was merely given data averaged across four very large groups of trials: left or right choices during slow or fast blocks. Nevertheless, the difference of activity related to left vs. right choices led the algorithm to assign a choice-related component that also happens to capture the evolving evidence for that choice. A neural population control method^84^ verified that this is not a simple consequence of tuning. Additional analyses (Extended Data Figure 6b) show that a unified evidence/choice component is obtained even if the PCA algorithm is only given data restricted to activity *after* movement onset. This suggests consistency between cell properties before versus after movement onset^54^, and argues against a categorical distinction between movement selection vs. execution circuits and in favor of a unified dynamical system, at least among the cells we recorded.

The emergence of the other components is also highly robust. If the PCA algorithm is only given data from the slow block, PC3 is lost (unsurprisingly), but the other components remain (Extended Data Figure 6c). Also not surprisingly, if we provide PCA with data from all 28 of our trial classes, the components are even more clearly distinguished, even though the number of cells that possess all of the required trials is reduced by 37% (Extended Data Figure 6d). In fact, any reasonable subset of data we have tried leads the PCA algorithm to identify the same functionally relevant components, albeit sometimes in a slightly different order depending on how much variance is captured by each. Finally, similar features are obtained if we leave out the cell duplication step (see Methods), although the resulting decision manifolds become asymmetrically distorted and some of the structure of loading matrices is more difficult to see (Extended Data Figure 7).

Finally, it is noteworthy that many of our findings reliably reproduce previous observations from very different tasks, including ones in which monkeys did not have to make any decision and were simply instructed to reach out to a single target. This includes the general flow of the neural state from target presentation to movement onset and offset^77^ (Figure 3a,c), the orderly relationship between reaction times and the neural state in PMd/M1^76^ (Figure 4a,b, purple ellipses), the compact subspace in M1 at movement onset^88^ (Figure 4a,b, green ellipses), and the presence of condition-independent components related to state transitions and elapsing time^79^ (Figure 2, PC1, PC4). Although most of our data comes from a prolonged period of deliberation that is not present in those other studies, the phenomena related to preparation and execution, which are shared between paradigms, are nevertheless robustly reproduced. This further strengthens the proposal that action selection and sensorimotor control are two modes of a single unified dynamical system.

Indeed, analyses of the loading matrix suggest that among the cells we recorded in PMd and M1, there is no categorical distinction between those involved in selection and those responsible for movement. Further analyses of synthetic populations (e.g. Extended Data Figure 8c) demonstrate that some of the higher order PCs we observed (PC5 and PC6, see Figure 2) may result from the heterogeneity of properties across our cell population, which includes purely decision-related and purely movement-related cells. However, analysis of the loading matrix concluded that these properties are not clustered into distinct categories as in synthetic data (Extended Data Figure 8d), but are instead distributed along a continuum (Figure 7).

While some of these findings could have been anticipated from analyses of individual cells, one important observation that would not have been possible concerns the shape of the decision subspace in the cortical populations. In particular, it is highly consistent across all of the different trial types and always resembles a thin manifold. This suggests strong normalization dynamics that are an inherent feature of recurrent attractor models. That is, the state of neural activity is pushed to lie on a surface that conserves some quantity (e.g. total neural activity with respect to some baseline) but is then free to move upon that surface under the influence of evidence, urgency, or simply noise. Furthermore, the particular shape of the manifold reveals consistent differences in the dynamics of different neural populations. In particular, regardless of what data we provide to PCA, the M1 manifold is always almost perfectly flat while the PMd manifold always exhibits a characteristic curvature. Interestingly, a very similar curvature was observed in preliminary analyses of PMd data in a very different decision task^110^. In contrast, the decision subspaces in the globus pallidus are much more compact and nothing like the thin manifolds in cortex.

We believe that the shape of the decision manifolds computed from different regions reveals properties of the underlying dynamics of neural activity in those regions. The planar manifold seen in M1 would be expected from a dynamical system with a steep interaction function governing mutual inhibition between competing groups of neurons (Figure 6c). In contrast, the curved PMd manifold would be expected from a system with a shallower interaction function (Figure 6e). In addition, we observed that the PMd manifold bends, approximately 200ms prior to commitment, along the component associated with the transition from deciding to acting. We suggest that this bend is the signature of a gradually emerging positive feedback in the cortico-striatal-thalamo-cortical circuit, which gradually overcomes inhibitory signals preventing premature selection^97^. Indeed, the analysis shown in Figure 5 reveals a strong correlation between how directional selectivity begins to emerge in GPi and how the PMd state begins to flow toward commitment, consistent with positive feedback between these regions^59,98,111^. Of course, establishing a causal relationship will require future studies, including simultaneous microstimulation in one region and recording in the other.

In conclusion, our analyses support the hypothesis that decisions between actions emerge as a competition within the sensorimotor system^7,9,25,91,102,104,105^, which is governed by recurrent attractor dynamics^23,24,26–28^. That competition is biased by sensory information coming at least in part from the prefrontal cortex^91^ and is gradually amplified by an urgency signal from the basal ganglia^44,108,109^. Commitment to a choice occurs through a positive feedback between premotor cortex and the basal ganglia^44,59,98,111^, leading to a winner-take-all process. That process then brings the cortical system to a state suitable for initiating the selected action^88,112^, setting into motion the “first cog” of a dynamical machine that controls our actions in the world^29–31^.

## Acknowledgements

This work was supported by Canadian Institutes of Health Research grants MOP-102662 and PJT-166014, the Canadian Foundation for Innovation, Fonds de Recherche en Santé du Québec, the EJLB Foundation to PC, and fellowships from the FYSSEN Foundation and the Groupe de Recherche sur le Système Nerveux Central to DT. The authors wish to thank Terrence Sanger for valuable observations about our data, and John Kalaska and Andrea Green for additional observations and helpful comments on the manuscript. PC and DT designed the study, DT collected the data; DT, JFC, AF, and PC analyzed the data; DT and PC wrote the manuscript; all authors approved the final version.

The data that support the findings of this study are available from the corresponding author upon reasonable request.

## Methods

### Subjects and apparatus

Neural recordings were performed in two male rhesus monkeys (*Macaca mulatta; monkey S*: 4-9 years old, 5-9kg; *monkey Z*: 4-6 years old, 4-7kg). Animals were implanted, under anesthesia and aseptic conditions, with a titanium head fixation post and recording chambers. The University of Montreal animal ethics committee approved surgery, testing procedure and animal care.

Monkeys sat head-fixed in a custom primate chair and performed two planar reaching tasks using a vertically oriented cordless stylus whose position was recorded by a digitizing tablet (*CalComp*, 125Hz). Their non-acting hand was restrained on an arm rest with Velcro bands. In most sessions, unconstrained eye movements were recorded using an infrared camera (*ASL*, 120Hz). Stimuli and continuous cursor feedback were projected onto a mirror suspended between the monkey’s gaze and the tablet, creating the illusion that they are in the plane of the tablet. Neural activity was recorded from the hemisphere contralateral to the acting hand with tungsten microelectrodes (1.0 – 1.5 MΩ, *Frederic Haer* and *Alpha-Omega Eng*.) moved with a computer-controlled microdrive (*NAN Instruments*) and electrophysiological signals were acquired with the AlphaLab system (*Alpha-Omega Eng*.). Spikes were sorted offline using Plexon Offline Sorter *(Plexon Inc.)*. Electrodes were targeted based on 3D reconstructions (Brainsight, Rogue Research) using structural MRI images (Siemens 3.0 T). Behavioral data were collected with the software used to display the task (LabView, *National Instruments*).

### Behavioral task and classification of trial types

Monkeys were trained to perform the “tokens” task (Figure 1a) in which they are presented with one central starting circle (1.75cm radius) and two peripheral target circles (1.75cm radius, arranged at 180° around a 5cm radius circle). The monkey begins each trial by placing a handle in the central circle, in which 15 small tokens are randomly arranged. The tokens then begin to jump, one-by-one every 200ms (“pre-decision interval”), from the center to one of the two peripheral targets always oriented at 180° to each other with respect to the center. The monkey’s task is to move the handle to the target that he believes will ultimately receive the majority of tokens. The monkey is allowed to make the decision as soon as he feels sufficiently confident, and has 500ms to bring the cursor into a target after leaving the center. When the monkey reaches a target, the remaining tokens move more quickly to their final targets (“post-decision interval”, which was either 150ms in “Slow” blocks or 50ms in “Fast” blocks (in a few sessions, the post-decision interval was reduced to 20ms in fast blocks). Once all tokens have jumped, visual feedback is provided to the monkey (the chosen target turns green for correct choices or red for error trials) and a drop of water or fruit juice is delivered for choosing the correct target. A 1500ms inter-trial interval precedes the following trial. We alternated between Slow and Fast blocks after about 75-125 trials, typically several times in each recording session.

The monkeys were also trained to perform a delayed reach (DR) task (usually 30-48 trials per recording session). In this task, the monkey again begins by placing the cursor in the central circle containing the 15 tokens. Next, one of six peripheral targets is presented (1.75cm radius, spaced at 60° intervals around a 5cm radius circle) and after a variable delay (500±100ms), the 15 tokens simultaneously jump into that target. This “GO signal” instructs the monkey to move the handle to the target to receive a drop of juice. This task is used to determine cell’s task response and tuning as well as the animal’s mean reaction time (RT), used as an estimate of the total delays attributable to sensory processing and response initiation. This quantity was then used to estimate the time of commitment in the tokens task. That is, the “decision time” (DT) in the tokens task was quantified as movement onset minus the mean RT from the DR task.

The tokens task allows us to calculate, at each moment in time, the “success probability” (SP) associated with choosing each target. To characterize the success probability profile for each trial, we calculated this quantity (with respect to the target ultimately chosen by the monkey) for each token jump (Figure 1b). For example, with a total of 15 tokens, if at a particular moment in time the right target contains *N_R_* tokens, the left contains *N_L_* tokens, and *N_C_* tokens remain in the center, then the probability that the target on the right will ultimately be the correct one (i.e., the success probability of guessing right) is:

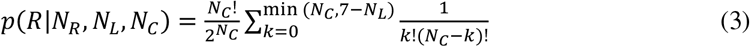

Although each token jump in every trial was completely random, we could classify *a posteriori* some specific classes of trials embedded in the fully random sequence (e.g. Figure 1c). In previous studies, we defined “easy”, “ambiguous”, and “misleading” trials on the basis of their success probability profile defined with respect to the first token jump. In contrast, because here we were primarily interested in examining activity with respect to commitment, we defined these trial types according to the success probability with respect to commitment time, estimated to be 280ms before movement onset. A trial was classified as “easy” if the SP was above 0.5 five tokens before commitment, above 0.55 three tokens before, and above 0.65 at the time of commitment. A trial was classified as “ambiguous” if SP was between 0.35 and 0.65 at five and three tokens before commitment as well as at the time of commitment. A trial was classified as “misleading” if SP was below 0.5 five tokens before commitment and then above 0.5 at the time of commitment. A trial was classified as “other” if it didn’t meet any of these criteria. Note that these four classes are non-overlapping. In the “Slow” block, we defined “early” trials as those where DT<1400 and “late” as those where DT≥1400ms. In the “Fast” block, “early” trials were defined as those where DT<950 and “late” as those where DT≥950ms.

In all tokens task trials, the targets were presented in opposite directions from the center, in two of six possible locations around the circle. Their placement was chosen according to the tuning of recorded cells, and in cases where a cell was not tuned, the leftmost and rightmost targets were used. Thus, in all cases there was a target to the left of the center and one to the right (sometimes at an oblique angle). Here, we grouped all of these into two groups: We defined the three targets between 90° and 270° as “left” targets, and the other three as “right” targets, and did not examine directional tuning in any more detail.

For the analyses in the present report, we defined 28 task conditions as follows. First, we separated trials into those recorded during the “Slow” block and those recorded during the “Fast” block, and in each block we split trials into those in which a “left” target was chosen by the monkey and those in which a “right” target was chosen. This yields 4 main groups: Slow-Left, Slow-Right, Fast-Left, and Fast-Right. Next, we split each of these four main groups into Easy, Ambiguous, Misleading, and Other, yielding another 16 conditions (Slow-Easy-Left, Slow-Easy-Right, Slow-Ambiguous-Left, etc.). Finally, we again split our four main groups into Early and Late, yielding another 8 conditions (Slow-Early-Left, Slow-Early-Right, Slow-Late-Left, etc.).

### Neural recording

Detailed methods for neural recording in PMd and M1 are described in Thura & Cisek^36^, and for recording in the globus pallidus in Thura & Cisek^44^. Recordings in dlPFC used the same methods as PMd/M1, and focused on the region just dorsal to the caudal end of the principal sulcus (Extended Data Figure 1). We used 2-4 independently moveable electrodes for cortical recordings, and one electrode at a time for recordings in the globus pallidus. Thus, most of the cells whose activity is analyzed and reported here were not recorded simultaneously.

In all sessions, we focused on cells showing any change of activity in the tokens task, and monkeys were usually performing the task while we were searching for cells. When one or more task related cells were isolated, we ran a block of 30-48 trials of the DR task to determine spatial tuning and select a preferred target (PT) for each cell (i.e. the target associated with the highest firing rate during one or more task epochs). Next, we ran blocks of tokens task trials using the PT of an isolated cell and the 180° opposite target (OT). We sometimes simultaneously recorded several task-related cells showing different spatial preferences, and since we always selected a single pair of targets, the actual best direction for all of the recorded cells was not always among these two. We usually started recording cells in the slow block because monkeys were more conservative in this condition. It was thus easier to assess cell properties online and more convenient to search for cells because fewer rewards were spent. When possible, cells were tested with multiple repetitions of slow and fast blocks to control for potential confounds related to evolving signals, elapsing time, and the monkey’s fatigue or satiation (see Thura and Cisek^43^ for control analyses on this question).

### Data analysis

A neuron was included in the analysis if it was recorded in both Slow and Fast blocks, thus including trials in each of the four main conditions: Slow-Left, Slow-Right, Fast-Left, and Fast-Right. This constraint was satisfied by 637 neurons out of the total 736 recorded across all brain regions (277 in PMd, 191 in M1, 52 in dlPFC, 41 in GPe, and 46 in GPi). For each neuron and each trial, neural activity was aligned to movement onset and the firing frequency was computed using partial spike intervals in ninety 20ms-bins from 1400ms before to 400ms after movement onset. The firing rate was then square root transformed and smoothed using a 25ms Gaussian kernel. Finally, we imposed symmetry on our neural population by duplicating each of our neurons with an identical “anti-neuron” that has the same activity in all trials, but with the trial labels switched between left and right choices. We describe our rationale for including this step below in the section “Rationale for duplicating neurons”.

To calculate the principal components of neural activity, we then grouped trials into our four main classes (Slow-Left, Slow-Right, Fast-Left, Fast-Right). In each group, neural activity was averaged together for each individual neuron regardless of the type of trial (easy, ambiguous, misleading, or other) and regardless of whether the choice was correct or not, or the reaction time was early or late. For each of these four classes, we constructed a 60×1274 matrix where rows are time bins (from 1000ms before to 200ms after movements onset) and columns are individual neurons. We then concatenated these four matrices into a 240×1274 matrix on which we performed standard Principal Component Analysis (using the *pca* function in Matlab 2019b), yielding a matrix of weights (“loading coefficients”) between cells and PCs as well as the variance explained by each PC. For further analysis, we only kept the top 20 PCs, which explained 97.9% of the total variance, though most of the interpretation will focus on the first four (total 80.3% of variance).

In addition to standard PCA, we also tried other dimensionality reduction methods, including factor analysis and Gaussian Process Factor Analysis (GPFA)^68,70^, but these yielded almost identical results (not shown). This is not surprising. For example, the major advantage of GPFA is that it jointly provides filtering with dimensionality reduction, which is critical when analyzing data from individual trials in which many neurons were recorded simultaneously. Our neurons were not recorded simultaneously, so we had to combine trials into similarity classes under the assumption that specific trial types were associated with similar neural activities on different occasions. This meant that we averaged activity across multiple similar trials, yielding much of the noise reduction that GPFA would have otherwise provided. Given the similarity of our results with different dimensionality reduction techniques, we chose to use PCA because it is the simplest, most widely known and well-understood, and ultimately easiest to interpret.

Using the 1274×20 loading matrix, we computed the average neural activity profile along each PC for a given trial condition and a given population of neurons by multiplying each neuron’s activity by its loading coefficient for that PC, summing all of these together, and dividing by the number of neurons in that population. We calculated confidence intervals around this average neural activity by randomly resampling trials, with replacement within a given trial condition, 1000 times for each neuron and time bin. We performed this process for all neurons together as well as for each of our neural populations separately (PMd, M1, dlPFC, GPe, and GPi).

Because not every neuron possessed a trial in each condition, potentially making it inappropriate to compare “PC space” trajectories from different conditions, we defined the following two groups of neurons designed to facilitate the analyses of primary interest. Group 1 consisted of neurons that possessed all conditions in the Slow block (Easy, Ambiguous, Misleading, Other, Early, and Late), for both Left and Right choices, as well as Early and Late conditions in the Fast block. This group was the focus of most of our analyses (e.g. Figure 3a,b; Figure 4; Figure 5), and included a total of 557 cells (258 in PMd, 173 in M1, 50 in dlPFC, 35 in GPe, and 41 in GPi). To facilitate comparisons between trial types in the Fast block (e.g. Figure 3c,d), we also defined Group 2, which consisted of neurons that possessed Easy, Ambiguous, Misleading, and Other conditions in the Fast block. This group included 452 cells (226 in PMd, 126 in M1, 46 in dlPFC, and 27 each in GPe and GPi). When calculating the temporal profile along the principal components (e.g. Figure 2), we used Group 1 neurons for all conditions except Easy, Ambiguous, Misleading, and Other trials in the Fast block. Although these definitions were used to make quantitative comparisons most accurate, they did not have a strong impact on the qualitative aspects of our data, which were similar even if we restricted all analyses to the 402 neurons that possessed trials in all 28 conditions (Extended Data Figure 6d).

Once we reconstructed the temporal profile along each PC for given conditions, we plotted each of them across time (Figure 2) and with respect to one another (e.g. Figure 3). To analyze the subspace visited by a neural population during the deliberation process, we examined PCs 1, 2, and 4 (see Figure 2) and constructed a 3-dimensional concave hull enclosing all of the neural states until the moment of commitment, for all conditions within a given block (using the *alphaShape* function in Matlab 2019b). For some of our brain regions this subspace resembles a thin 2-dimensional sheet, which we call the “decision manifold”. To quantify its thickness, we compared its surface area / volume ratio with that of a perfect sphere with equal volume, using the equation 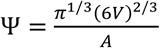, where A is the surface area and V is the volume^113^. The quantity Ψ is called “sphericity”, and is a dimensionless scalar that ranges from 0 for a 2D surface to 1 for a perfect sphere (the geometric shape with the smallest surface area / volume ratio). For the reader’s intuition, we note that an equal-sided cube of any size has Ψ=0.806 while a square sheet whose thickness is 1/50^th^ of the length of each side has Ψ=0.171.

To characterize the loading matrix, we considered each cell as a point in a high-dimensional space defined by its loading coefficients for the top 11 PCs (which accounted for 95.2% of the total variance). We then fit the resulting distribution of points for cells in a given brain region with Gaussian Mixture Models (GMMs) using the Expectation-Maximization algorithm (*fitgmdist* function in Matlab 2019b). For each fit, we tried GMMs consisting of anywhere from one to six 11-dimensional gaussians, performing 100 randomly initialized fits for each, and then used the Bayesian Information Criterion (BIC) to select the best fitting model. We found that with 100 randomly initialized fits, we reliably found the same best solution each time.

### Rationale for duplicating neurons

The rationale for duplicating the neurons stems from the fact that our population is inevitably a highly sparse under-sampling of the millions of neurons in the regions where we recorded. Importantly, we can assume that this under-sampling is not symmetric with respect to the proportions of neurons that contribute to movements to the left versus right. For example, we might have in our sample many high-firing cells preferring leftward movements, but fewer high-firing cells preferring rightward movements. Applying PCA to such asymmetrically under-sampled data will force the algorithm to try to capture this variance by aligning PCs to that asymmetry. We believe this is not informative. We can assume that our sampling is unlikely to be symmetric with respect to leftward or rightward movements, so it will not reflect any real asymmetry that may or may not exist in the brain. It will only distort our data in ways that will make interpretations more difficult.

For this reason, just before performing PCA we imposed symmetry on our data by using the “anti-neuron” approach classically used to produce population histograms of neural activity. In short, we assumed that for every given neuron that we recorded, there exists in the brain another neuron that we didn’t record, which has the same properties (same firing rate profile, same sensitivity to relevant variables, etc.) but has the opposite relationship with respect to the direction of movement. To implement this assumption, for each of our neurons we created a “sister” neuron with the same firing data except with the trial labels switched between left and right choices. We did this for all neurons regardless of whether they are tuned or not. All of the analyses shown in the main paper apply this anti-neuron duplication, but in the supplemental data we show analogous results obtained without that step. As can be seen (Extended Data Figure 7), the only difference is that the PCs produced without anti-neuron duplication are rotated and skewed versions of the ones produced after anti-neuron duplication, and their properties are slightly mixed between adjacent PCs. However, the general conclusions remain unchanged.

### Control analyses

To determine whether the PCs we found in the population data are a trivial consequence of properties of single neurons (e.g. directional tuning), we applied a control based on the Tensor Maximum Entropy (TME) method of Elsayed & Cunningham^84^, using code they provide at htpps://github.com/gamaleldin/TME. Briefly, this method preserves the primary first and second order covariance in the data along the temporal, neural, and condition-dependent dimensions but is otherwise maximally random. For example, tuning is preserved but not correlated across time. Any metric applied to analyzing the real data can then be compared to that same metric applied to surrogate data sets. If the metric computed from the real data lies within the distribution of that metric computed from surrogate data, then whatever is measured by the metric is a simple consequence of first or second order covariance.

Here, we were interested to know whether our finding of PCs that covary with evidence is a simple consequence of cell tuning. To that end, we computed the correlation between the evidence provided by token jumps (e.g. Figure 1c) and the temporal profiles of the components generated by PCA when applied to surrogate data. Our metric was equal to the maximum absolute value of the correlation coefficient for any of the top 10 components. We calculated this metric for the components produced using the real data (in which the best correlation was with PC2) and compared it to the distribution of the metric for 100 surrogate data sets generated using the TME method. That is, for each of the 100 surrogate data sets we performed PCA on the four main trial classes, then used these to calculate the temporal profile of the top 10 PCs for all trial types, correlated each of these with evidence, and used the highest correlation value. We consider the emergence of an evidence-related PC as a non-trivial consequence of tuning if the correlation metric of fewer than 5 of these 100 surrogate data sets is equal to or higher than the metric applied to the real data (i.e. p<0.05).

---

Extended Data Figures 1-8

Supplemental analysis: Distinguishing gradients versus clusters of cell properties

Supplemental references

**Extended Data Figure 1:**
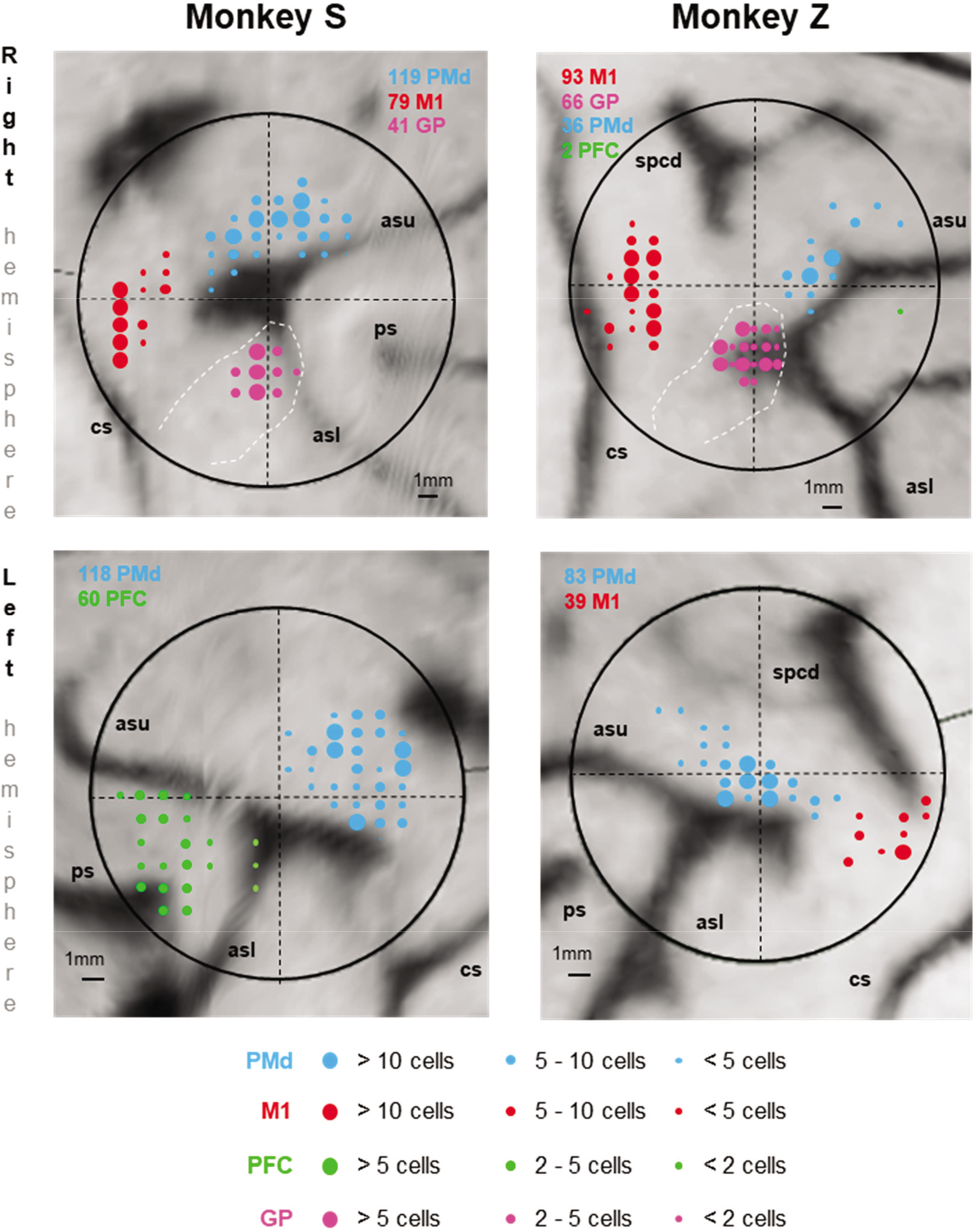
Recording locations. Colored circles indicate the penetration locations of recorded neurons, superimposed over the reconstruction of the brain surface using Brainsight (Rogue Research). Large black circles indicate the extent of the recording chamber. Medial is up and anterior is to the right in the top panels and to the left in the bottom panels. Abbreviations: cs – central sulcus; ps – principal sulcus; asu – upper limb of the arcuate sulcus; asl – lower limb of the arcuate sulcus; spcd – superior precentral dimple.

**Extended Data Figure 2:**
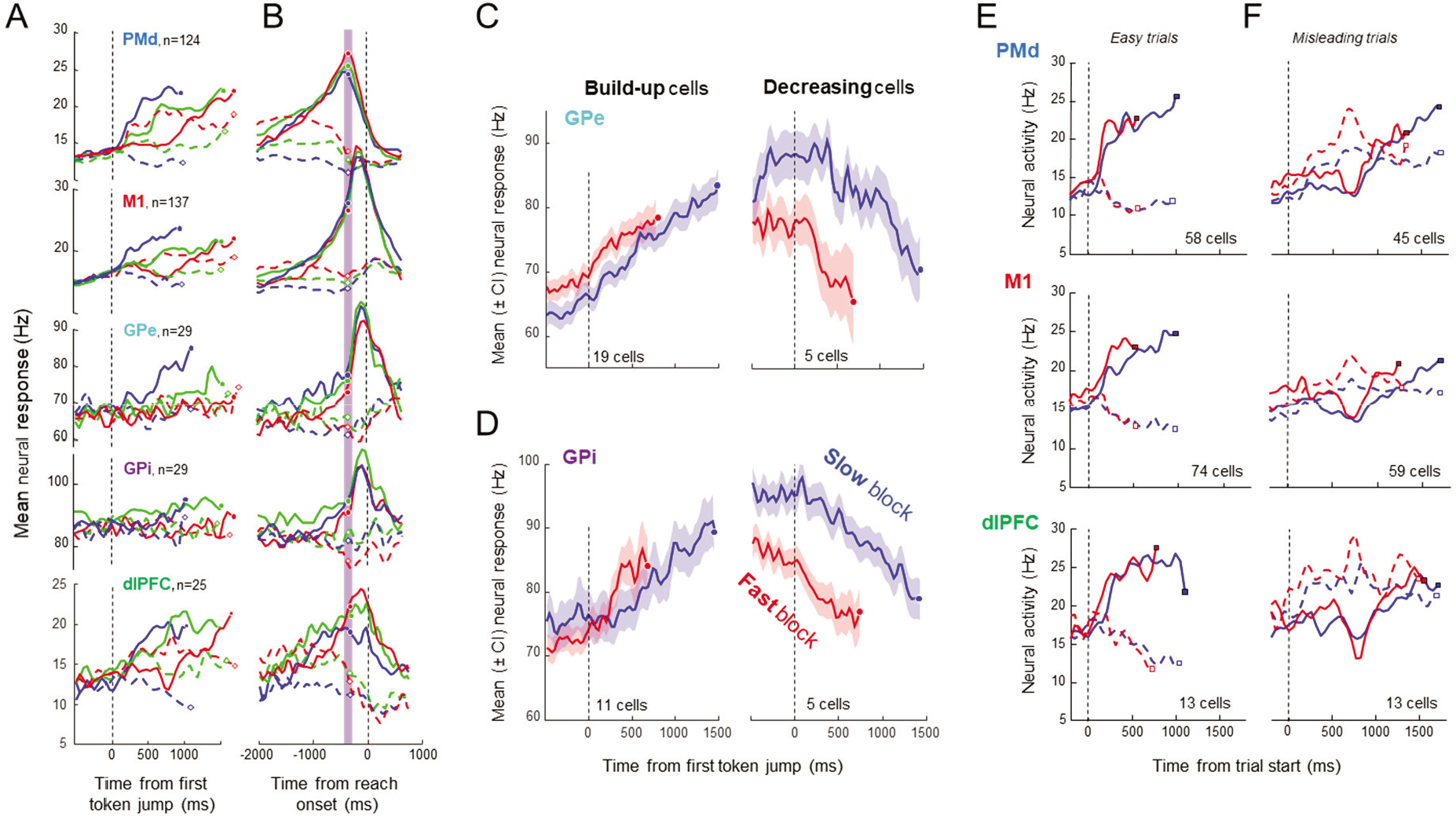
Neural activity during the tokens task. **A**. Average activity of directionally tuned neurons in five brain regions, computed separately for choices in each neuron’s preferred (solid) and opposite direction (dashed) for easy (blue), ambiguous (green), and misleading trials (red), during Slow blocks. Activity is aligned on the start of token jumps and truncated at decision commitment (circles), and only cells exhibiting tuning prior to movement onset are included. PMd: dorsal premotor cortex; M1: primary motor cortex; GPe: globus pallidus externus; GPi: globus pallidus internus; dlPFC: dorsolateral prefrontal cortex. **B**. Average activity of the same cells aligned on movement onset. The purple vertical bar indicates the estimated time of commitment, 280ms prior to movement onset. **C.**Average activity of build-up cells (left) and decreasing cells (right) in the GPe during Slow (blue) and Fast blocks (red), aligned on the first token jump. Shading indicates the 95% confidence interval. **D**. Same as C for cells in the GPi. **E**. Average neural activity in PMd, M1, and dlPFC during easy trials in Slow (blue) and Fast blocks (red), when the monkey chose the target in each neuron’s preferred (solid) or opposite direction (dashed). Here, only cells recorded in both blocks are included. **F**. Same as E for activity during misleading trials. Except for the PFC results, data in A-D is reproduced [with permission] from Thura & Cisek (2017) and E-F from Thura & Cisek (2016).

**Extended Data Figure 3:**
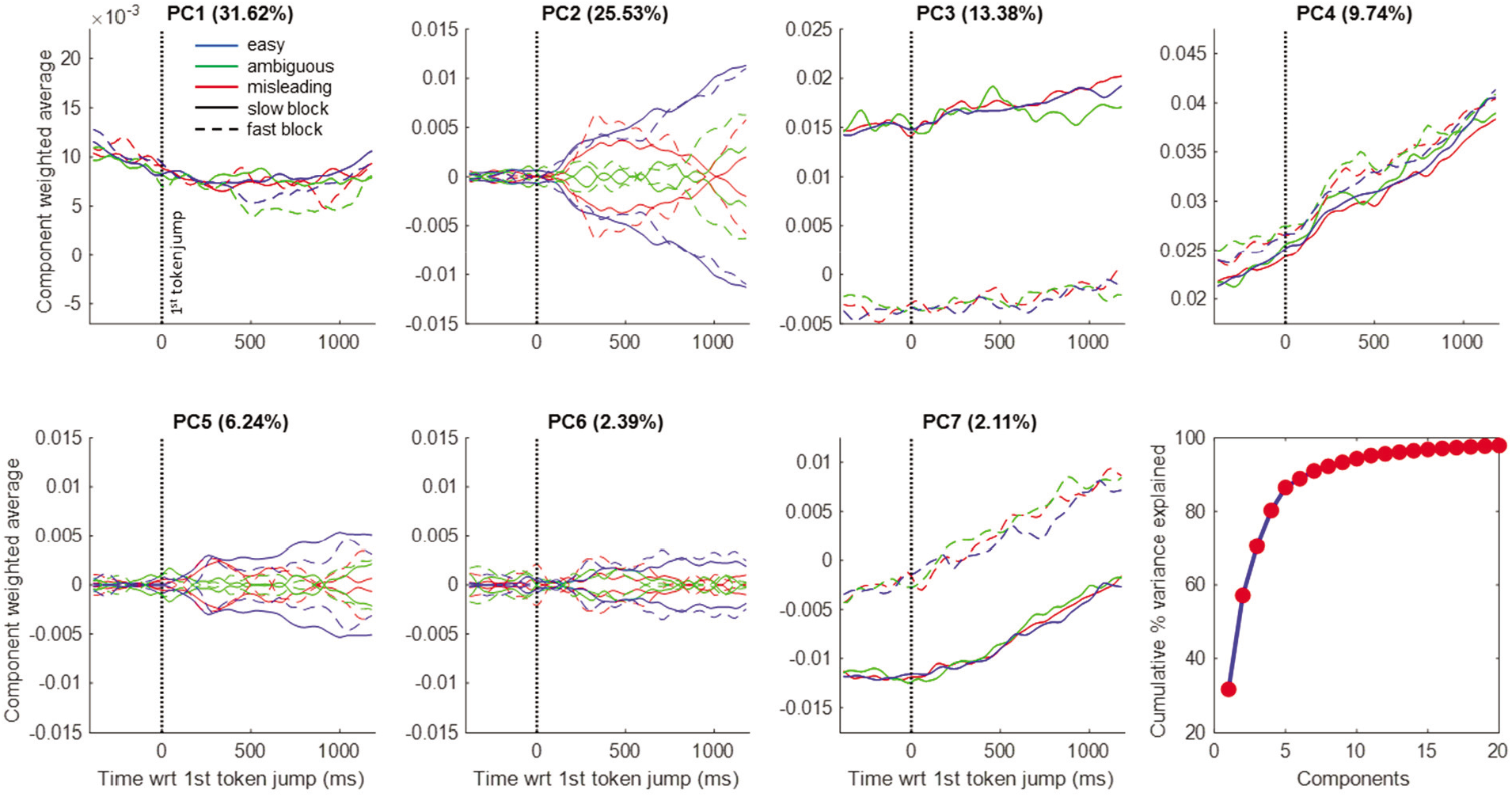
The temporal profile of the principal components (as in Figure 2) computed using data aligned on the first token jump (dotted vertical line).

**Extended Data Figure 4:**
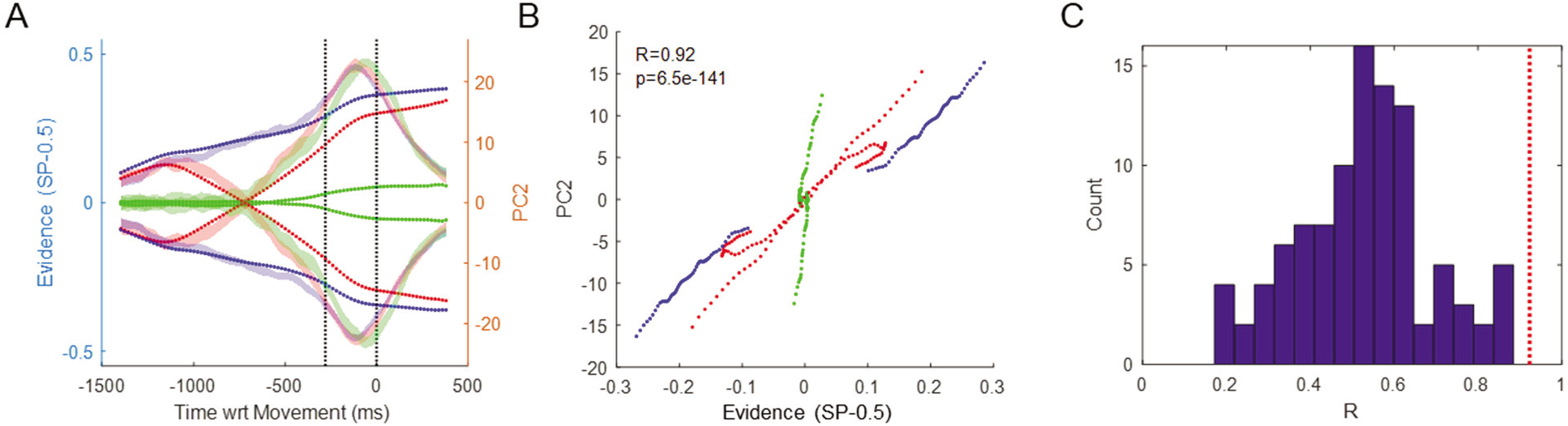
Comparison between PC2 and sensory evidence. **A.** Dotted lines show the sensory evidence in easy (blue), ambiguous (green), and misleading trials (red) calculated as the difference between success probability for the right target and 0.5. Shaded ribbons show the mean and 95% confidence interval of PC2 in the same trials (Slow block). The first vertical dotted line indicates commitment and the second indicates movement onset, on which all data is aligned. The evidence trace is delayed by 300ms, which provides the best fit. Note that until the moment of commitment, the pattern of PC2 closely resembles the evidence, except for diverging toward one of the choices even in the absence of evidence during ambiguous trials). **B.** The same data, *prior to commitment*, plotted as evidence versus PC2 in these six trial conditions. The correlation coefficient is R=0.9234 and p-value is well below 0.001. **C.** The distribution (N=100) of correlation coefficients obtained by performing the same analysis on surrogate data sets generated using the Tensor Maximum Entropy approach (see Methods). Here, each surrogate data set is represented as the highest correlation coefficient of any of the top 10 PCs against the profile of evidence. The mean R is 0.5294 (s.t.d.=0.1615). For comparison, the red line shows the R value from the real data (panel B), and it is higher than all of the R values from surrogate data (p<0.01).

**Extended Data Figure 5:**
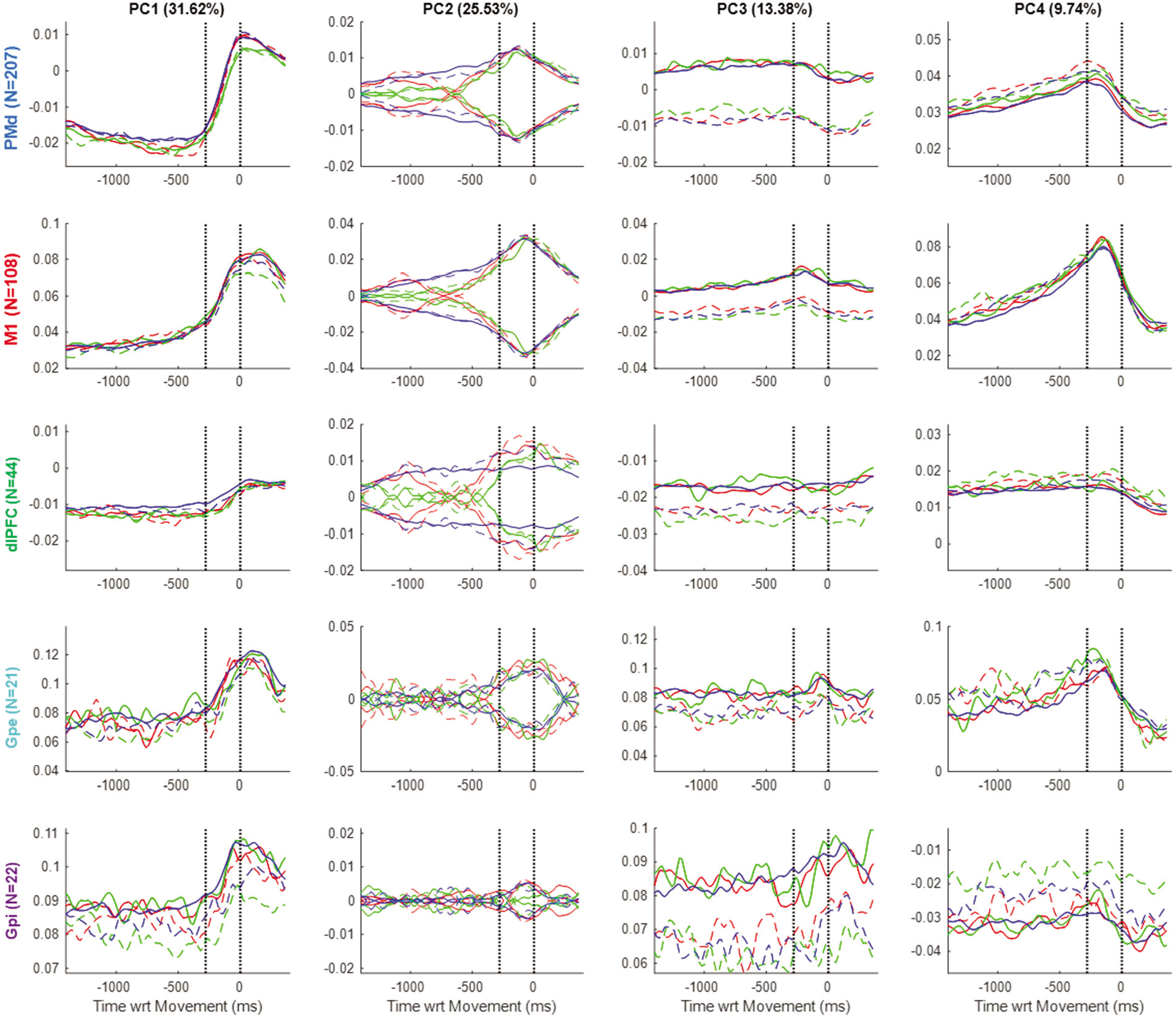
The top four principal components shown in Figure 2, but here computed separately from each of the five brain regions. Note: To facilitate comparison between Slow and Fast blocks, here we only include cells that were recorded in all trial conditions.

**Extended Data Figure 6:**
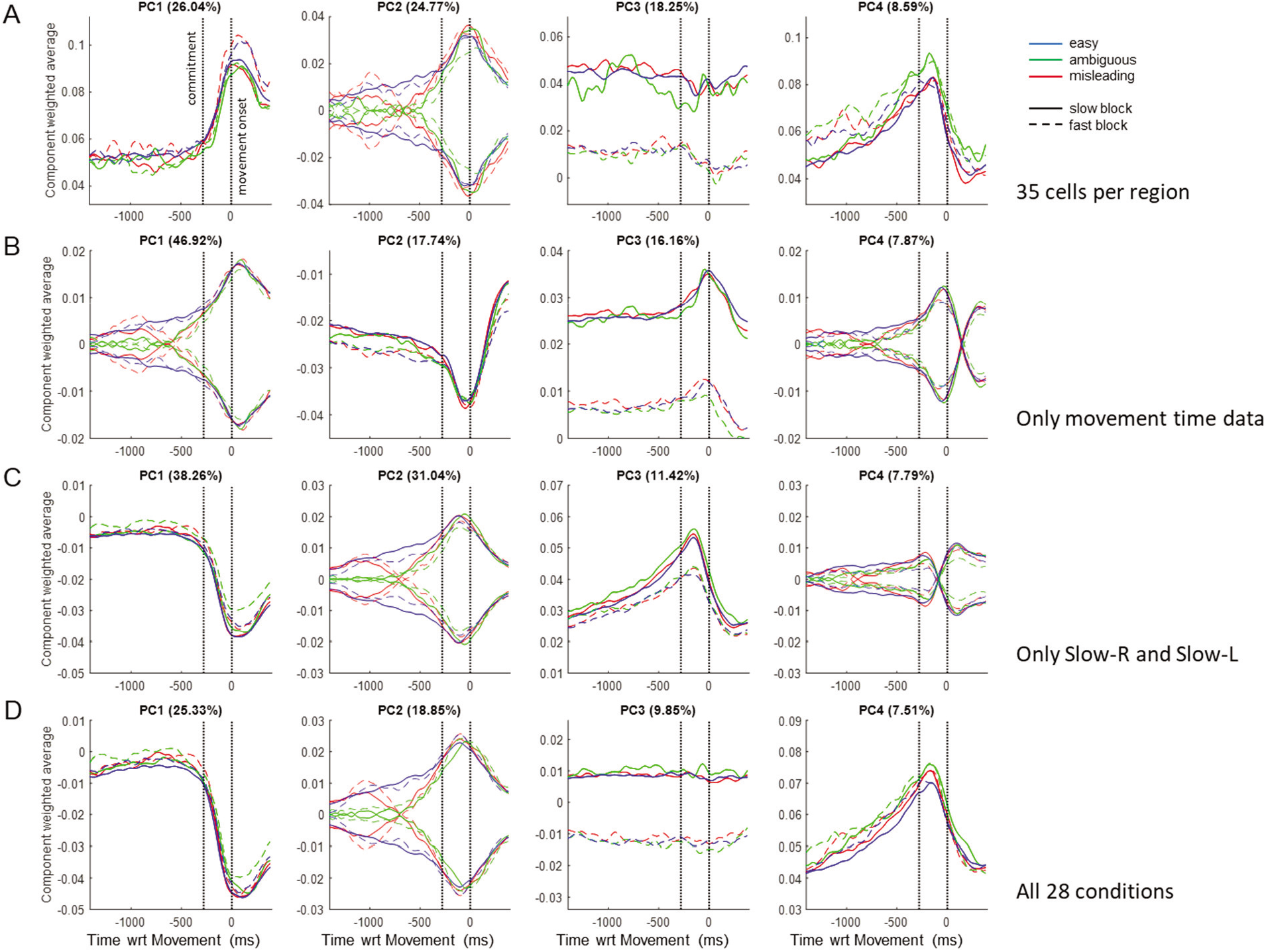
The first four principal components produced by providing PCA with different subsets of the data and then using the resulting loading matrix to compute PC profiles for all trials. **A.** PCs generated using data from only 35 neurons per region. Note that all of the general features of the PCs are similar to those in Figure 2, although they are more noisy. **B.** PCs generated using data only from the 400ms immediately following movement onset. The transition from deliberation to movement is now captured by PC2, the evidence by PC1, and the SAT by PC3. In contrast with the results reported in the main text, here there is no component related to elapsing time like PC4 in Figure 2. **C.** PCs generated using only data from Slow-Left and Slow-Right conditions. In comparison with Figure 2, there is now no SAT-related component and PC1 is inverted (note that the sign is arbitrary in PCA components). **D.** PCs generated using all 28 conditions, including activity from the 402 cells that possessed all of the 28 trial types. Note the similarity with the PCs shown in Figure 2, except for the inversion of PC1.

**Extended Data Figure 7:**
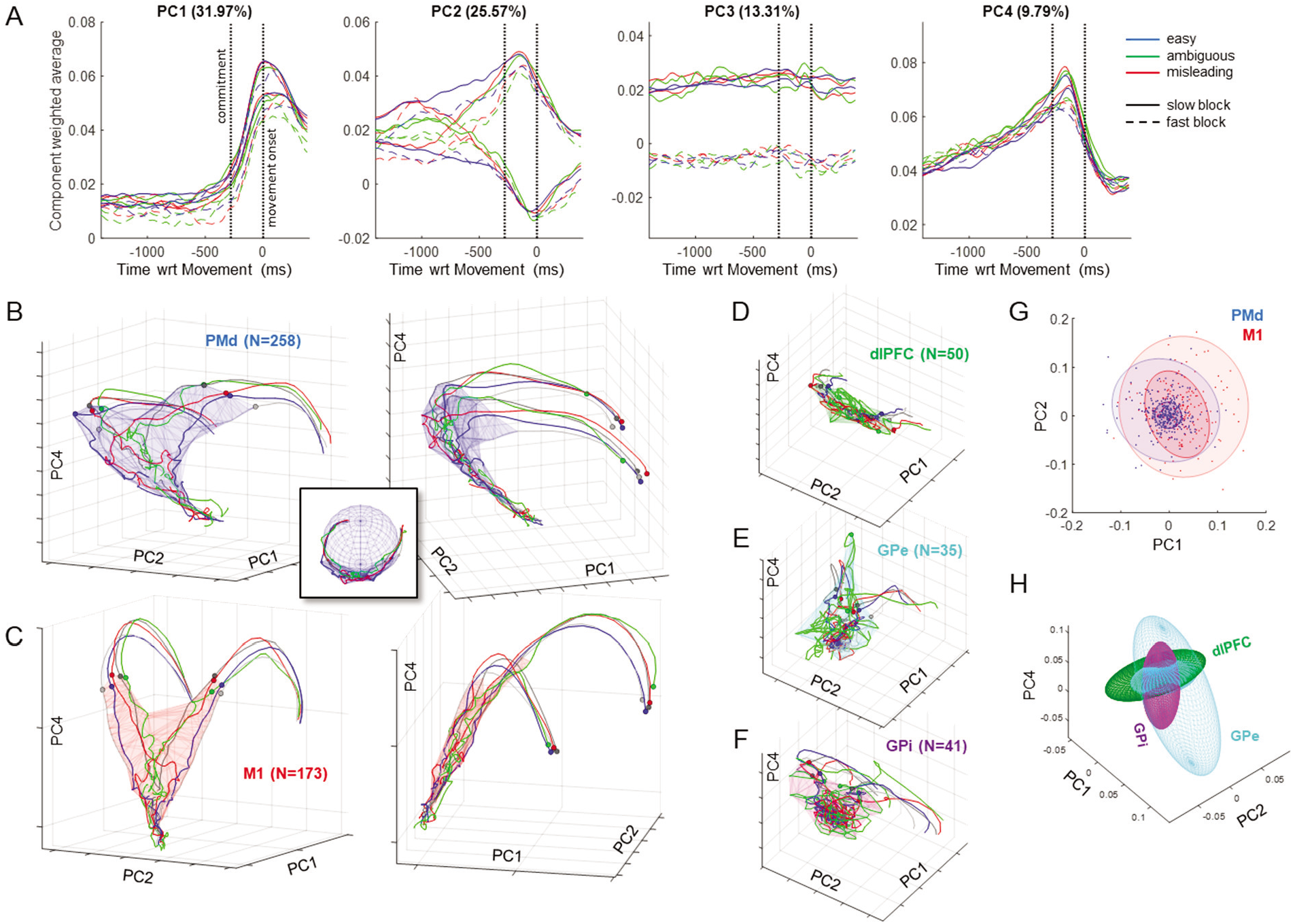
Results produced without the cell duplication step. A. The first four principal components. Note the similarity to the PCs shown in Figure 2, except without the imposed symmetry. Thus, PCs 1, 3 and 4 are now slightly different for the two choices, and PC 2 is no longer perfectly symmetric. B. Two views of the 3-D trajectories in PMd. Dots indicate the moment of commitment on the left panel, and movement initiation on the right panel. Note that the decision manifold (blue wireframe) is still curved as in Figure 4a, and relatively well fit by a sphere (inset). C. Two views of M1, same format as A. Note that the decision manifold (red wireframe) is still flat as in Figure 4b. D. A view of dlPFC. E. GPe. F. GPi. For all populations, the patterns are similar as data processed using the cell duplication step, except that symmetry is lost. G. Analysis of loading matrices for PMd (blue) and M1 (red). Both are still best fit by a mixture of two gaussians similar to those shown on Figure 7a-d, but their orientations are now rotated. H. The distribution of loadings in dlPFC (green), GPe (cyan), and GPi (purple). While the dlPFC distribution is still similar to that in Figure 7f, GPe is now best fit by a 2-gaussian mixture and GPi is rotated. Nevertheless, despite the rotations the distributions of dlPFC and GPi are still roughly orthogonal.

## Supplemental analysis: Distinguishing gradients versus clusters of cell properties

One of our key results is the apparent absence of distinct neural categories in our populations (Figure 7), arguing for a unified dynamical system of cells with continuous properties. However, is this continuity real or could it be an artifact of dimensionality reduction? To address this question, we created a variety of synthetic populations of neuron-like units, in which we deliberately created specific categories, and then applied to them the same analyses we used to examine real data. If our analyses fail to find these categories, then we cannot be confident that real categories do not exist in the real data. We attempted a variety of different ways to construct synthetic populations, but here report on two, each of which constructs synthetic units using combinations of the PCs we obtained from the real data (Figure 2).

The first population consisted of three distinct groups of cells. Group 1 consisted of 200 neurons with loading coefficients of 1 on PC1 and PC2, and zero for all others. Group 2 consisted of 200 neurons with loading of 1 on PC2 and PC4 and zero for all others. Group 3 consisted of 200 neurons with loading of 1 on PC1 and PC4 and zero on others. These loading coefficients were multiplied by 1+0.25*N*. where *N* is a normally distributed random variable with mean 0 and standard deviation 1. The synthetic activity of each cell was then computed as a sum of the temporal profile of each PC weighted by its loading coefficient for that cell. Finally, noise was added by multiplying the value of each bin of activity by a uniformly distributed random variable between 0 and 1.

We then subjected these synthetic units to exactly the same analyses we used for our real data, including square-root transformation, smoothing, and duplication. PCA analysis was then used to compute loading matrices of “synthetic” principal components (SPCs). The temporal profiles of the SPCs were then computed using a weighted sum of all units, and the loading matrix was analyzed using GMMs. For the synthetic data, we used 4-dimensional Gaussians to analyze the loading matrix.

As shown in Extended Data Figure 8a, the “synthetic” principal components (SPCs) almost perfectly capture the original PCs from which the cells were built, albeit in a different order (SPC1 is like PC2, SPC2 is like PC1, and SPC3 is like PC4). Furthermore, although the GMM analysis was applied to all 600 units together, without information on how the different groups were built, it correctly identified the relevant clusters (Extended Data Figure 8b).

The second synthetic population consisted of 600 neurons that all had loading coefficients of 1 on PC1 and PC2 and zero for others, again multiplied by 1+0.25*N*, turned into activity profiles using a weighted sum of the real PCs 1 and 2, and multiplied by a random variable uniformly distributed between 0 and 1. Next, these 600 units were split into three groups: For “deliberation” units, activity before commitment was multiplied by 1, activity between commitment and movement was scaled by a number linearly dropping from 1 to zero, and activity after movement onset was set to zero. For “movement” units, activity before commitment was set to zero, activity between commitment and movement was scaled by a number linearly rising from zero to 1, and activity after movement onset was multiplied by 1. For “mixed” units, all activity was kept unchanged.

We then ran PCA and obtained the synthetic components shown in Extended Data Figure 8c. Note that SPC1 is similar to PC1, SPC2 is like PC2, but now we also find an SPC3 that looks like PC2 except for a switch of activity between deliberation and movement. This is reminiscent of some of the higher components we found in the neural data (see PC5 and PC6 in Figure 2). Importantly, *no such component was used to build the synthetic units*. So where did it come from? The answer lies in the PCA algorithm, which sequentially identifies components on the basis of variance explained. After finding SPC1 it discovered SPC2, which together explain a large proportion of activity for both “deliberation” and “mixed” units. Since no linear combination of these two components could explain “movement” units, the next component (SPC3) captures some of the remaining unaccounted variance. In particular, note that a linear combination of SPC2 plus 3 times SPC3 can cancel out deliberation activity and produce a “movement” unit.

These observations on synthetic data suggest that the higher principal components we found in our real data (PCs 5, 6, 7, etc.) also do not necessarily reveal additional “higher order” features of neural dynamics, but simply result from the type of heterogeneity that has long been observed in sensorimotor cortical regions^1–6^. But this then revives the question of functional clusters of real neurons – might distinct categories exist in these regions? If they did, would our analysis of loading matrices identify them correctly? Extended Data Figure 8d shows the result of GMM fits to the entire population of 600 synthetic units, again performed without any information on the underlying groups. Clearly, the relevant clusters were found. In particular, “movement” units (red) were identified with two Gaussians (one for right-tuned and one for left-tuned units), loaded positively onto SPC1, and onto both SPC2 and SPC3 with 3 times larger coefficients for the latter – consequently cancelling out their deliberation-time activity. In contrast, “deliberation” units (blue) were identified with two oriented Gaussians orthogonal to those of “movement” units and negatively loaded on SPC1. Finally, “mixed” units (green) were identified as clusters lying in-between. This demonstrates that if distinct categories of neurons really did exist in the real populations we recorded, the GMM analysis would have identified them correctly.

**Extended Data Figure 8.**
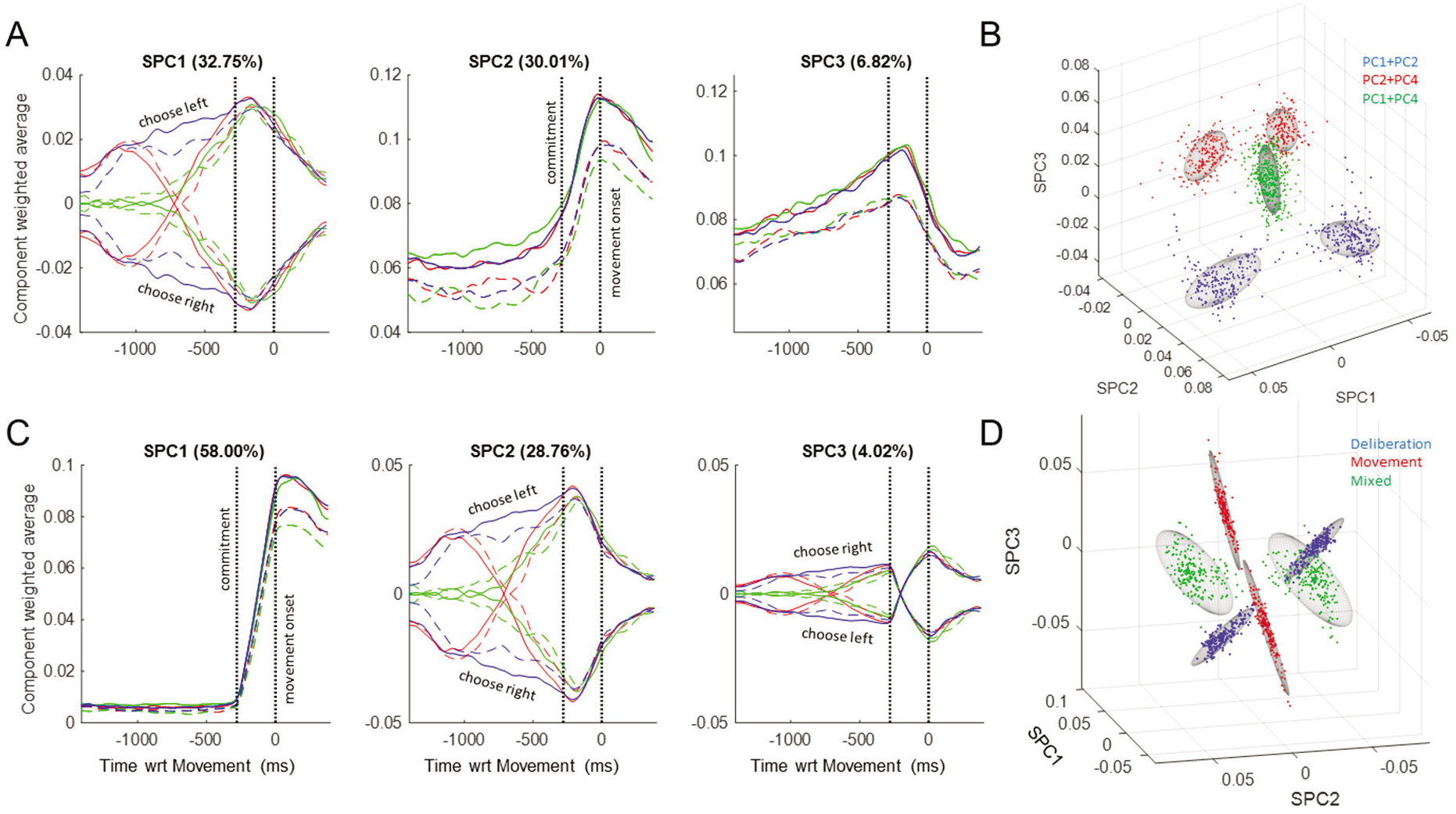
Analyses of synthetic populations. **A**. The top 3 Synthetic PCs (SPCs) obtained from a population of 600 neurons constructed using combinations of the original PCs 1, 2, and 4 (see Figure 2). **B**. The loading coefficients of all 600 neurons, fitted with GMMs (gray ellipsoids). Colors indicate the three groups of neurons built with different combinations of those original PCs (see legend). **C**. The top 3 SPCs obtained from a population of 600 neurons constructed only using the original PC1 and PC2, but separated into three distinct categories: cells that are only active during deliberation, cells only active during movement, and cells active during both epochs. **D**. The loading coefficients of all 600 neurons, fitted with GMMs (gray ellipsoids). Colors indicate the three categories of neurons.

## Notes

### Competing Interest Statement

The authors have declared no competing interest.

### Summary of Updates

Acknowledgements section updated

